# Simple syntactic rules through rapid synaptic changes

**DOI:** 10.1101/2023.12.21.572018

**Authors:** Lin Sun, Sanjay G Manohar

## Abstract

Syntax is a central organizing component of human language but few models explain how it may be implemented in neurons. We combined two rapid synaptic rules to demonstrate how neurons can implement a simple grammar without accounting for the hierarchical property of syntax. Words bind to syntactic roles (e.g. “dog” as subject or object) and the roles obey ordering rules (e.g. subject → verb → object), guided by predefined syntactic knowledge. We find that, like humans, the model recalls sentences better than shuffled word-lists, and when given the permitted role orderings, and a set of words, the model can select a grammatical ordering and serialize the words to form a sentence influenced by the priming effect (e.g. producing a sentence in the passive voice after input of a different sentence also in the passive voice). The model also supports languages reliant on affixes, rather than word order, to define grammatical roles, exhibits syntactic priming and demonstrates typical patterns of aphasia when damaged. Crucially, it achieves these using an intuitive representation where words fill roles, allowing structured cognition.

## Introduction

Why is it easier to remember “colorless green ideas sleep furiously”, than “furiously sleep ideas green colorless”? Although the sequence of words is itself new, something about the order of word *types* is familiar. In a grammatically correct sentence, each word has a role, which makes it more likely to be followed by a word with another particular role. This underlying syntactic representation is key both in language comprehension (Jones & Farrell, 2018; Perham et al., 2009), and production (Kathryn Bock, 1986; Pickering & Ferreira, 2008). However, we do not yet have an understanding of how the brain encodes syntax in real-time as words are presented in a string via our senses. Understanding this sequential nature of syntax may itself be an important mechanism for reasoning and high-level cognition (Süß et al., 2002; Verguts & De Boeck, 2002).

Currently, no dynamic neural model can capture the distinction between the *structure* and *content* of thoughts. Specifically, how do brain circuits capture the fact that when we think, our sequence of thoughts has a structure that obeys a set of rules, while the contents are highly flexible?

Sequences can be understood as forward associations between items (Wennekers et al., 2006), but in the case of syntax, it is the word-*roles*, not words themselves, that have sequential structure. We hypothesize that working memory links each word to its role in the sentence, and those roles follow probabilistic orderings. Although this cannot account for true grammar in the study of linguistics since it does not support hierarchical structure (Everaert et al., 2015; Poletiek et al., 2021), it provides a biological perspective for understanding some aspects of syntax, as words, and their associated roles, are processed in the order of our sensory input. In this paper we propose a biologically plausible neural architecture that stores and generates syntactic sequences.

Many neural models of working memory have been proposed to hold sequential information and they have adopted a range of methods to simulate syntax and structure within sequences such as storing either timing (Tokuhara et al., 2021), order (Rolls & Deco, 2015), or production rules (Cer & O’Reilly, 2006; Kruijne et al., 2021; O’Reilly & Frank, 2006). Word sequences may form forward associations between words (Rezende et al., 2011; Zhang et al., 2022) with short-term synaptic plasticity (Kappel et al., 2014) or by associating each word with a “temporal context” (neurons that represent the current time) in a “phonological loop” (Howard & Kahana, 2002; O’Reilly & Soto, 2001). But these models ignore the rules that assign words to their structural roles. They do *not* account for the effect of syntactic rules, e.g. that word lists are recalled better if they are syntactic than non-syntactic.

To account for syntax, patterns of neural activity can encode sentences (Kriete et al., 2013; Markert et al., 2005) through word-and-role combinations (Rolls & Deco, 2015). However this quickly leads to a combinatorial problem with longer sentences, e.g. requiring duplicate neurons to allow encoding “dog” as either subject or object. Similarly, tensor-product representations bind words to roles but also require units coding every possible filler in every possible role (Smolensky, 1990). One option has been to interface neural networks with an external symbolic system (Hammond & Leake, 2023). A more linguistically inspired strategy is to build a syntactic tree as words are encoded (Hagoort, 2003; Reitter et al., 2011), however these models do not specify how words are stored together with the structure.

Fully neural systems such as transformer-based large language models use backpropagation to train weights in a deep neural network. Transformer-based large language models, in comparison to symbolic approaches, have superior generative power and can easily fit large data sets. However, deep-learning models do not include explicit filler-role binding, and while they have demonstrated some limited abilities in discerning grammatical and ungrammatical sentences (Warstadt et al., 2019), their training process has little resemblance to real-world language acquisition. They also encounter difficulties when role-position information is critical (Clouatre et al., 2022), and rely on shifting buffers which are difficult to implement using biological neurons.

Recurrent neural networks (RNN) trained on next word prediction may be more biologically plausible and can represent rich language structures (Gu & Lim, 2022; Marvin & Linzen, 2018; Schrimpf et al., 2021; Suzgun et al., 2019; Tokuhara et al., 2021). However these networks still lack an intuitively symbol-like architecture (Dehaene et al., 2022).

To address this, alternative approaches have used RNNs to predict syntax by training them to generate grammar parsing *actions* (e.g. “open new noun phrase”, “close phrase”) (Brennan et al., 2020; Kuncoro et al., 2016). While these RNN grammars represent syntactic information, they do not store the resulting structured sentence. Other RNNs have incorporated separate attention mechanisms for syntactic and semantic information (Russin et al., 2019). This mechanism can generalize grammatical rules to new content words, but requires tricks like reading sentences both forward and backward in parallel, and it is unclear whether the RNN keeps a representation of the whole sentence.

In summary, previous neural models of language tend to either fail in biological plausibility, or fail to separate form from content in their internal representations. This is a binding problem: a general issue which applies across human cognition (Roskies, 1999). While potential neural solutions have been proposed (Manohar et al., 2019), they have not been tested for language.

We propose that words and their roles are coded by two populations of neurons. Each role within a sequence is associatively bound to a different word – or equivalently, each word is tagged with a different syntactic role. While neurons in superior temporal cortex are selective for the identity of auditory stimuli (Belin et al., 2000), neurons in prefrontal cortex encode abstract categorisation of input (Antzoulatos & Miller, 2014; Freedman et al., 2003; Shima et al., 2007; Tanji et al., 2007) and might therefore represent word roles (Gwilliams et al., 2022). We implement this role tagging with rapid synaptic changes that can store *bindings* between words (“fillers”) and their roles. Since synapses vastly outnumber neurons, the combinatorial binding problem is solved.

We harness an idea first proposed for working memory, where contents are rapidly bound to ‘slots’ using a population of competing, flexible neurons (Manohar et al., 2019). Here in the current model, words are bound to roles in a similar way. We use a highly simplified, hand-wired network to simulate hearing sentences, and also producing them from unordered words.

### High-level Overview of our Model

Our approach is to first tackle the minimal case, using just a handful of neurons, with hand-wired synapses and rapid plasticity. There is no training or training data, since the connections are so simple, and are pre-specified for a given language. Word-selective neurons and role-selective neurons form a recurrent network (Figure 1a). The two types of neuron are bidirectionally connected, and role neurons also have directed connections to each other (Figure A1). To flexibly encode a new sentence, word-selective neurons are activated in sequential order. These words drive role-selective neurons according to long-term knowledge of word classes encoded in synapses between word and role neurons. A role may be filled by one of several words (e.g. a subject noun could be “dog” or “cat”), and a given word could play different roles (e.g. “dog” could be a subject noun or object noun). In addition to this drive from words, role neurons drive each other. Long-term knowledge between role neurons provides a tendency to follow a familiar order (e.g. subject → verb → object) forming a Markov chain. Together these two constraints determine the role associated with each word.

**Figure 1.**
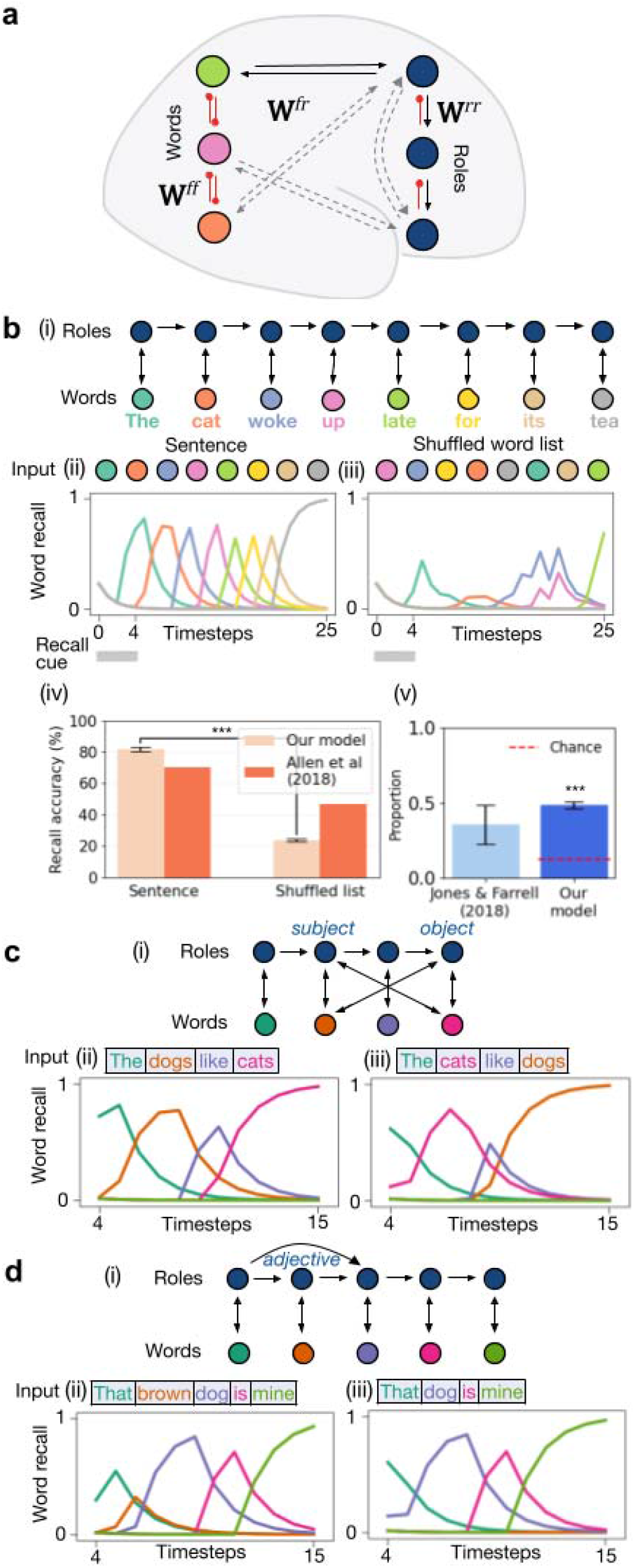
Simulations of simple grammar with our model. **a**, Architecture of our model comprising word and role-selective neurons in a rapidly reconfigurable neural network. The word neurons compete with each other via fixed interactions (**W**^***ff***^), but connect reciprocally to and from the role neurons (**W**^***fr***^). The role neurons also compete amongst themselves (**W**^***rr***^), with some role sequences being prepotent. b, Simulation 1: Recall in a case of simplest possible grammar ((i) no branching). The model receives a recall cue to generate the input - either a (ii) syntactic sentence or a (iii) shuffled word-list. (iv) The model demonstrates better recall accuracy for a sentence compared to a shuffled word-list, comparable to human data from Allen et al (2018). (v) In fact, recall of the shuffled word-list tends to produce syntactic word pairs (“grammaticalization errors”) above chance, comparable to human data from Jones & Farrell (2018). c, Simulation 2: (i) The network can sequentially encode two sentences with the same syntax but different words. Both nouns “dogs” and “cats” are able to bind to either the subject or object role neurons. (ii) Recall of the first sentence input where “dogs” is the subject and “cats” is the object. (iii) Subsequent recall of the second sentence, where “dogs” and “cats” switched positions. d, Simulation 3: (i) The network can encode sentences with different syntactic structures. (ii) The network encodes and recalls a first sentence with an adjective, and then (iii) a sentence without an adjective. *p<0.05; **p<0.01; ***p < 0.001, error bars are standard error of mean (SEM). Black arrows: connections with predefined long-term knowledge. Red rounded arrow: inhibitory connections.

Superimposed on the long-term synapses are short-term, rapidly changing weights. These weights between the word and its role, and between a role and its successor, are rapidly strengthened via a Hebbian rule, encoding the sequence of word-role pairs into working memory. The temporarily strengthened connections generate a *plastic attractor* – a partly-stable sequence of states of the network, that allows the subsequent re-activation (i.e. recall) of the same sequence of words and roles. Our model runs in continuous time and is robust to the timing of inputs, maintaining information across delays using these partly-stable states.

### Base Model Technical Details

Here we introduce the technical details of the base model. Additional key details, if necessary, outlining each specific simulation will be included in their respective sections. Comprehensive outline of each individual simulation is also included in Supplementary Methods.

The two main neuron classes in our base model are words, i.e. fillers (**f**), and roles (**r**) (Figure 1a). They follow simple firing dynamics and are connected by synaptic weights. Long term background weights (**W**_**L**_) encode long-term knowledge about language. Short-term weights (**W**_**S**_) indicate transient changes in synaptic strength, and are added on top of the long-term weights, to store information for a given sentence as it is heard or generated. They may correspond to changes in calcium or phosphorylation in post-synaptic terminals.

Long-term weights from word to role 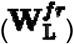 encode knowledge about which roles a word can fill. Long-term weights from role to role 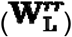 encode which roles can follow which others. Short-term word-to-role weights 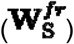 store the words that fill each role in the current sentence. Short-term role-to-role weights 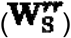 retain the structure, i.e. role order, of the current sentence.

To encode a sentence, words are encoded as a one-hot vector **f** and are sequentially activated by auditory inputs, and drive role neurons **r** (Figure A1a). The word-role synapses 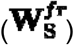 are strengthened bidirectionally. At the same time, the activation of a new role suppresses other roles, by winner-takes-all dynamics, but also rapidly strengthens synapses from the previously active role to the newly active role 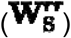 (Figure A1b). These two kinds of rapid plasticity respectively retain a working memory of the words that fill each role, and which roles follow which. So, when the sequence is later triggered by driving the first role neuron, the corresponding word neurons are re-activated in sequence (Figure A1c).

We term this encoding and retrieval behavior a ‘plastic attractor’, since the input words create a set of partly-stable states, each state corresponding to a role paired with a word. Syntax constrains the role dynamics, so the sequence of states tends to align with a grammatical sequence of words.

### Connectivity

Neurons have three main sources of input: external, feedback from neurons of other types, and dynamics among neurons of the same type (Equation 1).

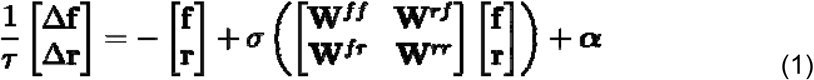

Neurons receive external activation **α** through sensory input. Word neurons **f** can drive role neurons **r** via **W**^***fr***^ **= (W**^***fr***^)^T^. Words mutually inhibit each other by a fixed amount **W**^***ff***^ **= − 1.5 · 1**(i.e. all elements of the matrix), and role neurons are connected to each other with a structured sequential pattern given by **W**^**rr**^. Activity is limited by a sigmoid activation function ***σ*** described below in Equation 7. Activity decays over time with rate −1. The constant ***τ* = 0.5** determines the rate of updates.

Sensory input during memory encoding comprises a one-hot sequence of words, and then at the very start of recall, one role neuron receives a short burst of direct input as trigger.

Firing rates **f, r** are truncated to lie between 0 and 1, 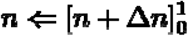.

Weights **W**^***fr***^ and **W**^**rr**^ are the sum of a stable long-term knowledge component **W**_**L**_ and rapidly plastic short-term component **W**_**S**_, in addition to noise (***e***) (uniform distribution) where applicable (Equation 2).

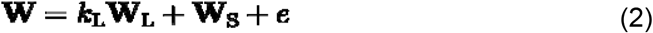

Long-term weights **W**_**L**_ are scaled from 0 to 1, so that ***k***_**L**_ is the maximum strength of prior knowledge. The rapidly changing components of the weights, **W**_**S**_, is updated based on Hebbian rule (Equation 3).

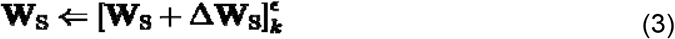

The square bracket indicates the value is truncated to values between ***k*** and ***ϵ*** (Equation 4).

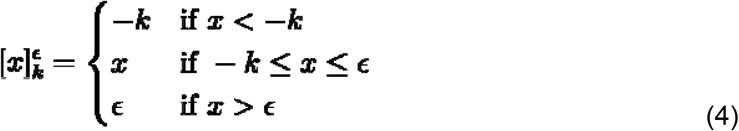

Overall this means that **W**_**S**_ varies from **−*k*** to **+ ϵ. W**_**L**_ is fixed as a pattern of either 0 or 1 and pre-specified in all simulations to reflect prior understanding of word classes and role order.

### Plasticity rules

There are key differences between Hebbian rules in word-to-role connections and in role-to-role connections. 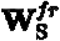 is strengthened when a role neuron and word neuron are simultaneously active, but weakened if only one is active (Equation 5). ***h*** is a constant that controls the balance between strengthening and weakening, and **λ**^***fr***^ **= 1** is the learning rate.

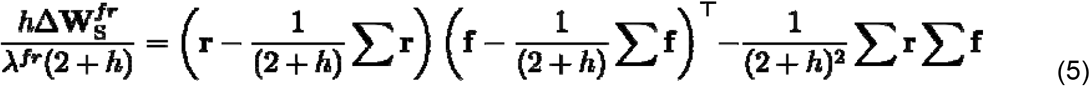

For 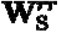, the connection weight from role neurons ***r***_**1**_ to ***r***_**2**_ is strengthened during rapid encoding if both role neurons are simultaneously active and paired with a high level of input from the word neurons to ***r***_**2**_ (Equation 6). In other words, if a role neuron is being driven by an external word, then it will learn to receive stronger input from other roles that were also recently active, but those synapses will weaken otherwise. The diagonal entries of **W**^***rr***^ is kept constant at zero and **λ**^***rr***^ is the learning rate. **⊙** is element wise multiplication.

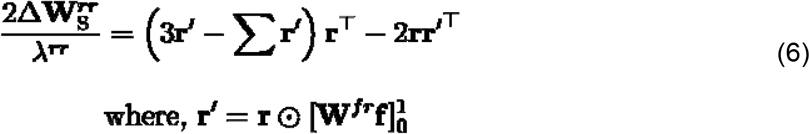

### Steps and Constants

Words are presented sequentially with no gaps. Each period consists of one run of the complete input (i.e. a sentence or a word-list). Recall phase is prompted by activating the first role neuron for five timesteps, followed by no further direct input. Analysed result begins from the fifth timestep onwards. The neurons are expected to spontaneously retrieve the input with help from a leftward shift of the sigmoid function to increase all neuronal tendency to self-activate. The equations of the recall phase are identical to the encoding phase except that **W**_**S**_ is not updated.

Other than these Hebbian updates, there is no built in weight decay.

Equation 7 is a sigmoid softmax function with slope 10, where ***θ*** is 0.5 during all encoding and zero during all recall.

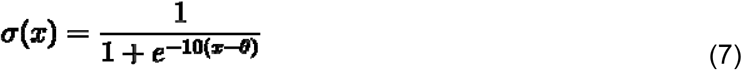

Table A1 summarizes constants unspecified above. Graphical representations of Equations 5 and 6 are presented in Figure A2. The pseudocode for the algorithm of each timestep is presented in Figure A3. All statistical analyses were performed with Scipy 1.5.0. Python code and random initialisations were created with random seeds of Python 3.8.5 and NumPy 1.19.1.

### Computer Code

Jupyter Notebooks for our experiments are available at https://doi.org/10.5281/zenodo.10372949.

### Simulating Simple Grammar in Working Memory

We begin with the simplest possible syntax using only our base model, and extend it incrementally to evaluate the model’s core capabilities.

First we simulate a grammar with only one possible syntax, and where only one word can fulfill each role (Figure 1b). As expected, the model recalls syntactic sentences with greater accuracy than a shuffled word-list. The pre-existing long-term syntactic knowledge means that role neurons are activated in order. However with a shuffled list, the role neurons receive two conflicting drives: the word neurons activate their corresponding roles, but this does not match the natural tendency of the role-to-role synapses to generate a grammatical sequence. Consequently, errors occur, resembling human data (Allen et al., 2018). Furthermore, when humans recall shuffled-order sentences, they make “grammaticalization” errors where adjacent words are more likely to fit with grammar than chance (Jones & Farrell, 2018). Our model gives rise to this because role neurons tend to activate in grammatical sequences.

Next we show that the model can store which word is bound to each role (Figure 1c). Two word neurons can compete to drive one role neuron, for example either “dogs” or “cats” could be the subject. Similarly, a single word could fill one of two possible roles, e.g. “dogs” may play either the subject or object role in the sentence. To remember which word fills which role, the word-to-role synapses rapidly strengthen to create a new attractor state, potentially overwriting other connections of those units.

Finally, we demonstrate that the network can remember which of two syntactic structures was presented, by rapid plasticity between the role neurons (Figure 1d). For example, the chain of role neurons can branch as adjectives can be skipped. The network remembers which particular sentence structure was used by strengthening the role-to-role synapses. This channels the sequence of role neurons during recall. If motifs from both Simulations 2 and 3 are combined, the network simultaneously remembers both the words’ roles and the branches taken (Figure A4a).

Since the model allows multiple role sequences and multiple roles for a word, ambiguity naturally arises. The long-term strengths of word-to-role synapses determine the probability of a word being used in a particular role, and simultaneously, the long-term strengths of role-to-role synapses determine the transition probabilities to the next role, given the history. The competition results in parsing of sentences with soft constraint satisfaction, as observed in humans (Cho et al., 2017) (Figure A4b). The disambiguation of word class based on role-sequence is also supported by evidence from magneto-encephalography studies (Gwilliams et al., 2023) (Figure A4c).

### Rationale for Directional Hebbian Rule

Directional plasticity is a key innovation of our proposed network. When the role-to-role directional Hebbian rule is replaced with a standard Hebbian rule defined between word and role neurons (Figure A5a), the network fails to capture the role sequence despite successfully binding words to their roles, and therefore fails to encode the simple linear sentence in Simulation 1.

Why is standard Hebbian plasticity insufficient? Hebbian encoding between two sequential role neurons occurs only when both role neurons are activated, which happens during the first role neuron’s decay and the second role neuron’s ascent. During this small window, the first role neuron loses activation from the previous word neuron while the second role neuron is being activated by the input from the next word neuron. Since both role neurons are active, standard Hebbian plasticity would strengthen both forwards and backwards connections. Our directional plasticity captures the fact that this *difference in input determines which direction* to strengthen the role-to-role connection in, to build the overall role sequence.

One alternative to uni-directional plasticity would be to force the network to run in a forward direction only, by using very strong long-term role-to-role weights. However, when role-to-role connections become sufficiently strong to achieve this, they also drive activation of all role neurons following the present role neuron binding to the current word input. This disrupts the binding of one word to only one role neuron (Figure A5b).

### Winner-takes-all Dynamic

Strong inhibitory connections create competition that results in the selection of the most strongly activated neuron of each class to be activated while the other neurons in the class are silenced.

For roles, the mutual inhibition is what generates the ‘slot-like’ representation of the components in the sentence. By removing this inhibition (Figure A5c) for connections between role and role neurons, all role neurons downstream of an activated role neuron in a pre-defined syntactic sequence are activated simultaneously as a result of long-term role-to-role weights. Therefore, word neurons fail to bind to the appropriate role neuron due to the presence of several simultaneously activated role neurons.

Furthermore, the rate at which the preceding role neuron decays is amplified by permitting inhibition between role and word neurons. This minimizes the time window where both the preceding role neuron and the following word neuron are activated simultaneously. Without this, spurious associations can form between the preceding role and the following word neurons.

On the other hand, if the inhibition is too strong, it will cause a reverse effect where the preceding word neuron suppresses the activation of the following role neuron. Therefore, the winner-take-all dynamics in word neurons does not solely depend on the inhibition between word and role neurons. We include an additional inhibitory fixed connection between the word neurons (i.e <u>**W**^***ff***^</u>) (Figure A5d).

### Languages where affixes, rather than word order, define grammatical roles

In many languages, word order is not fixed. Instead, affixes attached to word stems provide cues to a word’s role. Ultimately the roles assigned to words depend on a combination of constraints including word order and affixes. To model this, the word neurons function as stems (e.g. “lik-” in liking) allowing them to fill a range of roles depending on affixes (e.g. “-ing”, “-able”) (Anderson et al., 2016). In other languages such as Latin, nouns have different affixes for their nominative and accusative cases. By using word neurons as stems (e.g. “can-”) and affix neurons as suffixes (e.g. “-is” or “-em”), we avoid duplicating every noun by allowing suffixes to flexibly pair with different stems.

It turns out that no extra machinery is needed for our network to model this. We create affix neurons that function like word neurons, but are distinct from words as they need to be concurrently active without mutually inhibiting word neurons (e.g. “can-” is not competing with “-is”). Each affix neuron is paired with a role neuron via a long-term, fixed connection (e.g. “-is” and the subject role neuron) (Figure 2a, bold solid arrows).

**Figure 2.**
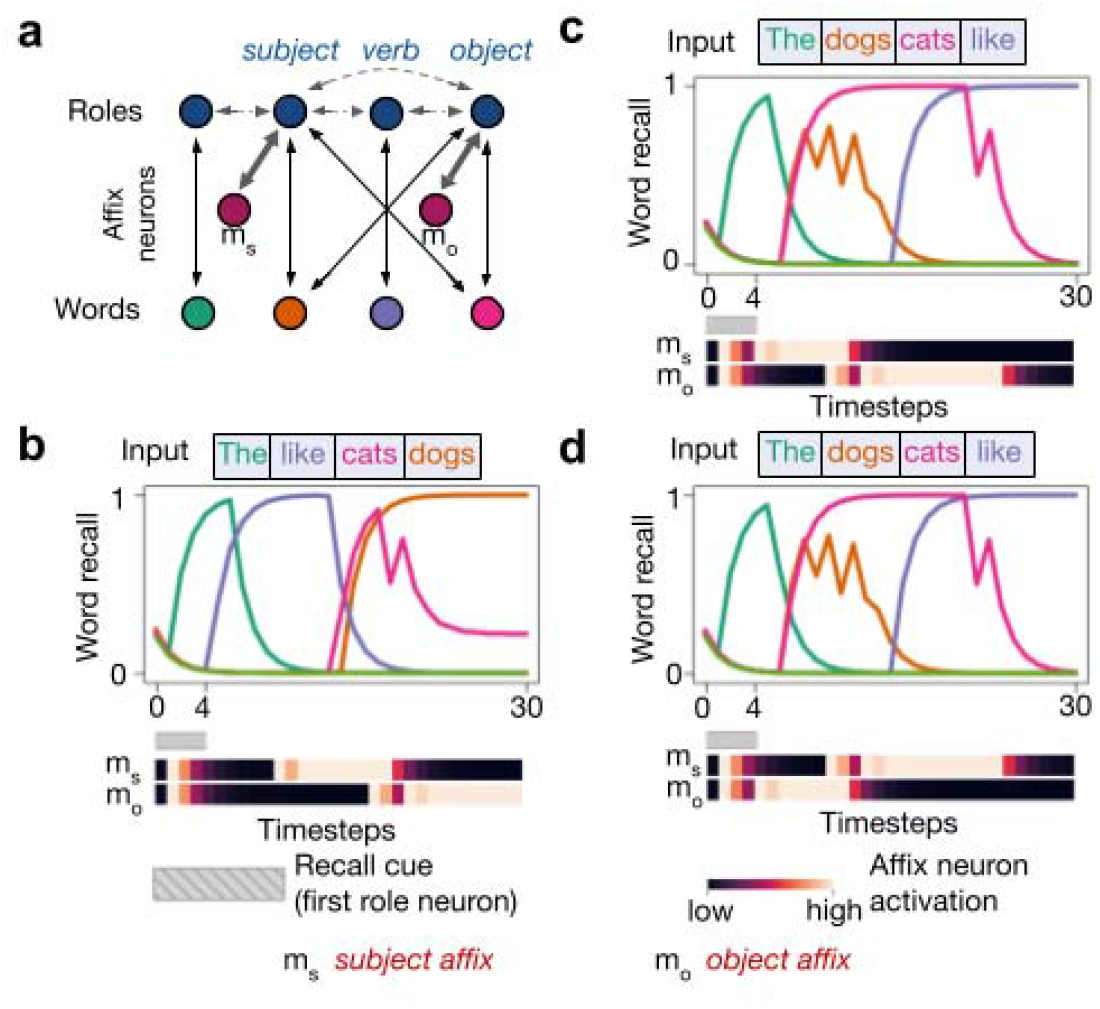
Affixes cue word roles. **a**, In this network, affixes accompany word input in a language with no long-term knowledge of syntactic structure. Each role neuron has no bias towards any single role neuron (broken arrows, not all shown). Therefore, the order in which the words are presented during the encoding phase will determine the syntactic structure during recall. Bold solid arrows: fixed strong connection between pairs of affix and role neurons. b, Simulation 4: Encoding and recall of an input sentence with the structure “verb-subject-object”. c, Simulation 5: Same for “subject-object-verb”. d, Simulation 6: Same for “object-subject-verb” but with the nouns in the same serial position as c. Note the similar activation in word neurons but different activation sequences in affix neurons.

During encoding, if a word requires an affix to determine its role, we activate both the word *and* affix neurons simultaneously. The affix dictates the syntactic roles attached to each word (overriding word order effects) by directly driving the corresponding role neuron.

### Affix Neurons

Concretely, affix neurons (**m**) operate like words (Equation 8), with the differences that there is no lateral inhibition between words and affixes, and they have stronger drive to the roles:

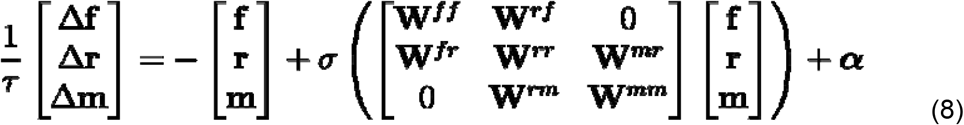

where **m** is a one-hot vector representing a currently active affix,**W**^***mm***^ **= −10.1** (mutual inhibition between affixes) aid selection between the affixes, and **W**^***rm***^ cause affixes to drive a specific role (e.g. subject or object). Affix neurons receive and return activation to its paired role neurons with **W**^***rm***^ = **W**^***mr***^(direct activation between affix and respective role neuron) being fixed non-encodable constants.

### Affix Simulations

In simulations 4-6, the network encodes and recalls sentences in a language with no constraint on word order in the long-term role-to-role synapses (Figure 2b-d). The word order is instead encoded by the sequence of affix activation during encoding. Due to rapid changes in the role-to-role synapses, the network recalls words in the same order they were encoded despite lacking long-term word order knowledge. The affix neurons have mutual inhibition separate from the word neurons, so that appropriate word-affix pairs will activate simultaneously during encoding to ensure the relevant grammatical role is activated with the word (e.g. “dogs” + affix neuron for subject when “dogs” is in the subject position of the sentence). Across languages, our model can flexibly accommodate a wide range of possible structures, from supporting affix to order-based, including situations where affixes are paired only to some words.

### Sentence Generation: Syntactic Serialization

A central feature of human cognition is that we can express ideas. In particular, we arrange words into ordered sentences, to express an idea held in an internal semantic representation. We demonstrate that our model is able to serialize a “bag of words” that are fed simultaneously to the network (Figure 3a,b). Words receiving input compete with each other, but the order in which they become active during recall is constrained by role-to-role synapses. The model even inserts function words that were not present in the input (Figure 3c). To achieve this, a ‘conceptual’ input must be added to the word neurons (Technical change to Equation 1 outlined below), and role-to-role weights must be stronger to promote automatic sequential word activation. We will in a later section demonstrate that this sequentialization is also influenced by syntactic priming.

**Figure 3.**
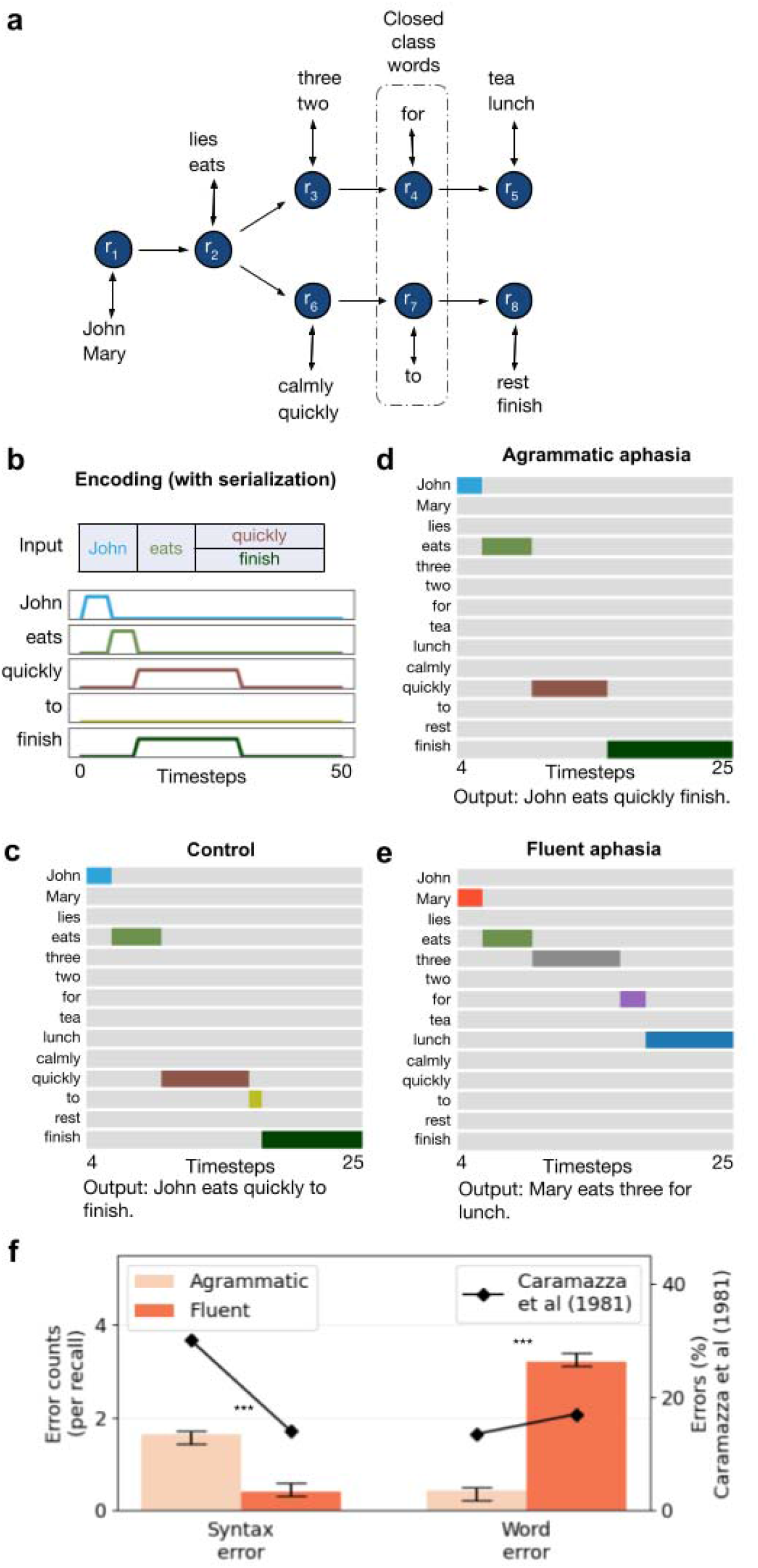
Sentence generation and aphasia **a**, Network architecture for simulating syntactic serialization and agrammatic and fluent aphasias. b, During the encoding phase, “quickly” and “finish” are fed into the network simultaneously without the closed-class words. c, Simulation 7: The network serializes the word inputs, adding the missing closed-class word, to form a syntactic sentence. d, Simulation 8: The agrammatic lesion results in omission of the closed-class word while the message is preserved. Coloured maps of word activation represent the strongest activated word of each timestep. e, Simulation 9: In fluent aphasia, grammatical structure is preserved but there are word substitution errors. f, Frequency of syntax errors versus word errors in agrammatic and fluent aphasias produced by the model (Simulations 8 and 9), compared to data from humans. Agrammatic and fluent aphasia simulations include noise, therefore more example outputs are presented in Figure A6.

### A Technical Detail of Serialization

A key technical difference between simulations with serialization and the base case simulation is that ***α*** in Equation 1 is included within the sigmoid function for Simulations 7, 8, 9 and 12 (Simulation 12 is part of the later section - syntactic priming). Shifting ***α*** into the sigmoid function allows role-to-role encoding guided solely by long-term syntactic knowledge without need for explicit order in the word input. This can apply to syntactic serialization of non-auditory input.

### Like humans, lesions to the model produce dissociable deficits in structure vs. content

Neurological disorders can disrupt language production. Strikingly, some patients lose grammar without losing content (agrammatic aphasia) whereas others lose contents of speech with preserved structure (syndromes of fluent aphasias). A key characteristic of agrammatism, often seen in Broca’s aphasia, is a breakdown in sentence structure despite the correct words being generated (Bates et al., 1988). In particular, their speech production is characteristically non-fluent with frequent pauses, and telegraphic speech with missing closed-class function words (Friederici, 1982). Despite this, content words are relatively preserved. So for example, an agrammatic patient might utter, “John… um… finish,” instead of “John eats quickly to finish”.

### Agrammatic Aphasia

Since we model only one neuron per role, to simulate lesions we added noise, corresponding to weaker representational strength. To model agrammatic aphasia, we introduced noise into rapid synaptic plasticity between role neurons. We further assumed that certain roles can only be filled by a limited set of words (closed-class words, e.g. prepositions). The model produces errors characteristic of agrammatic aphasia (Figure 3d) (Friederici, 1982; Martínez-Ferreiro et al., 2019).

### Fluent Aphasia

In contrast to the non-fluent output from our simulation of agrammatic aphasia, we are able to reproduce fluent aphasia characterised by semantic word substitution: choosing the wrong word, but with preserved grammar. Semantic word substitution is a feature that manifests across several aphasia subtypes, including jargon aphasia (Hillis et al., 1999; Marshall, 2006), paragrammatic aphasia (Maviş et al., 2020) and semantic aphasia. These types of aphasia also often feature poor comprehension, preserved fluency of speech, and poor insight. We model the preserved grammar in fluent aphasia by adding noise to the long-term word-to-role knowledge with rapid synaptic plasticities between word and role neurons switched off (Figure 3e). We compare these results qualitatively with empirical observations (Figure 3f) (Caramazza et al., 1981). In the case of fluent aphasia, we compare our output to empirical data from paragrammatic aphasia.

### Lexical and Syntactic Priming

Synaptic changes also provide immediate explanations for both lexical and syntactic priming. In the model, lexical priming occurs when exposure to the same item improves retrieval (Tulving & Schacter, 1990) and syntactic priming improves recall accuracy when target and priming sentences have the same syntax (Figure 4a) (Bock et al., 1992; Bock & Griffin, 2000; Kathryn Bock, 1986; Pickering & Ferreira, 2008). The network also takes longer to repeat a sentence when the syntax changes (F. Chang et al., 2006; Jaeger & Snider, 2013; Malhotra, 2009; Reitter et al., 2011). Intuitively, this is because encoding a sentence creates stable attractor states, which make the network less effective at shifting to new states, yielding priming for both words in their roles, and also sequences of roles (Figure 4a).

**Figure 4.**
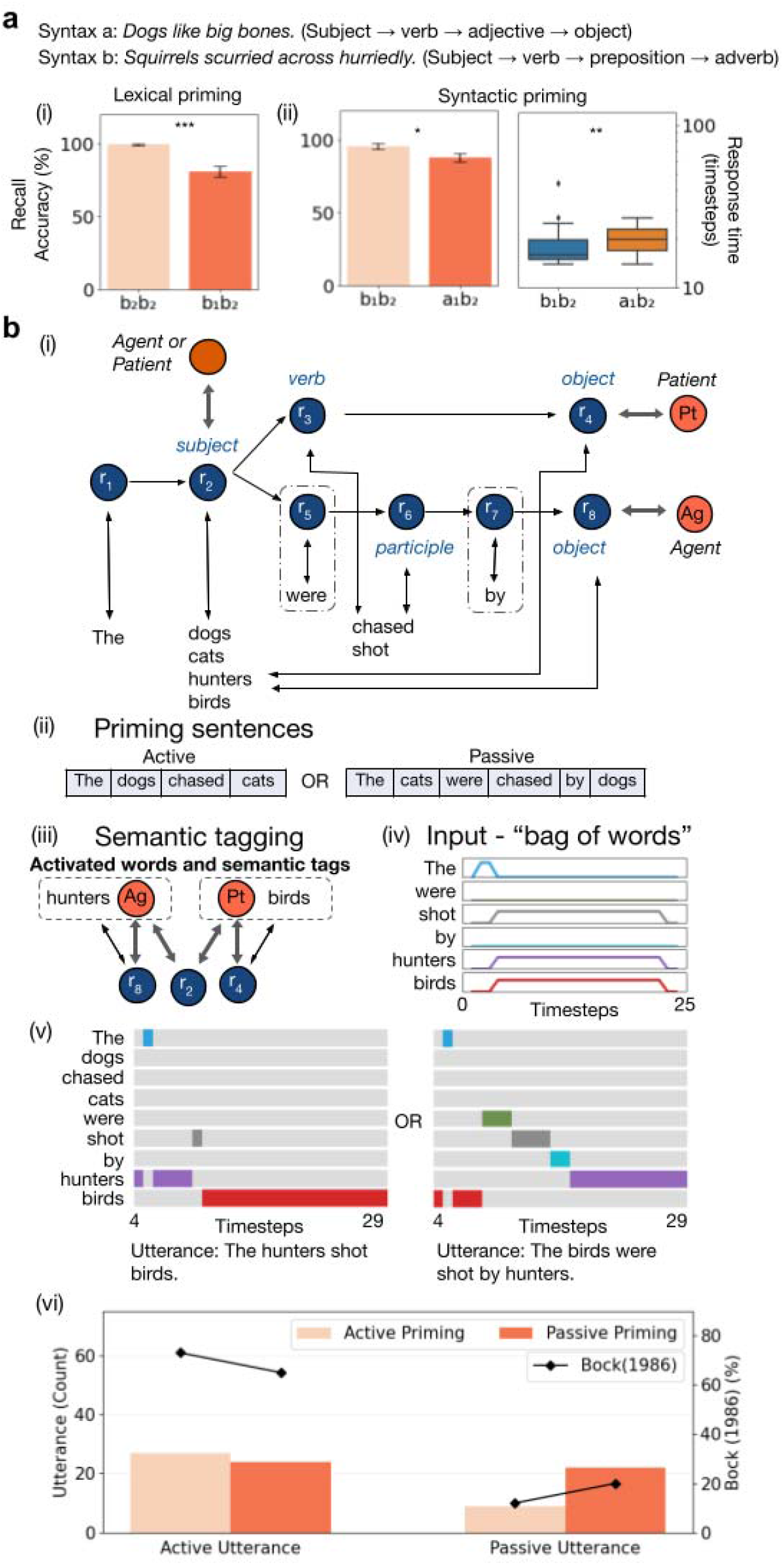
Synaptic changes predict priming **a**, Lexical and syntactic priming during comprehension. (i) Simulation 10: Lexical priming - recall accuracy improves on repeated encoding of the same sentence. (ii) Simulation 11: Syntactic priming - recall accuracy is reduced following encoding of a syntactically distinct sentence. Response time is also increased. Different syntactic structures are coded with letters, a or b. Numerical subscripts (a_1_, a_2_ etc) represent different sentences with non-overlapping words. More lexical and syntactic priming results are presented in Figure A7. ns non-significant; *P-value<0.05; **P-value<0.01; ***P-value < 0.001, error bars are SEM. Box plot shows interquartile range. b, Simulation 12: Syntactic priming during production. (i) To encode active and passive sentences, the network has two branching pathways through the role neurons. There are two object role neurons, each with a semantic tag (one for agent, and the other, patient). The subject can either be an agent or patient. (ii) The network is first presented with an active or passive priming sentence. (iii) Then on encountering a mental idea, both nouns are semantically associated with the relevant agent or patient tag. (iv) “Bag of words” without closed-class words (i.e. “were” and “by”) were fed simultaneously to the network to allow syntactic serialization. (v) The network generates either the active or passive voice. (vi) The syntactic priming effect qualitatively matches human data.

Syntactic priming can also bias language production (Figure 4b) (Kathryn Bock, 1986). The network is first primed with a sentence in either the active or passive voice. Then, the content elements in a scenario are pre-associated to their appropriate semantic roles (i.e. nouns to ‘agent’ or ‘patient’) via semantic tags. The semantic tags behave just like the affix neurons, preferentially driving different role neurons. Finally, the words necessary for forming sentences are simultaneously fed to the model as a “bag of words”. The roles activate sequentially with a preference for the primed order, generating words in the order appropriate for that syntactic construction. This reproduces the empirical finding that a syntactic prime biases language production.

Semantic tags can also guide the flow of role neurons, for example allowing the phenomenon of “affix hopping” where semantic tags can control affixes after the word to which they apply (Figure A8).

So far, we assumed that long-term word-class and role-to-role knowledge are already pre-learned and that plastic attractors are created *within* the existing knowledge structure.

Next we challenge the model to acquire syntactic knowledge using slow plasticity, by simply adding long term plasticity that mirrors short term changes.

To simulate a naive learner, the long-term synaptic weights are initially empty, both from words to roles (encoding which roles a word can fill) and from role to role (encoding which orders of roles are permitted). Whereas previously these long-term weights were specified by hand and were fixed, now they can vary. They have learning rates that are much lower than for the short-term weights, but still follow the same Hebbian rules.

### Long-term learning rule

The long term weight matrix **W**_**L**_ is gradually updated with Equation 9 in small steps to simulate long-term learning.

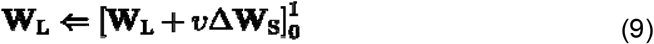

Here,***v* = 0.001** for long-term word class knowledge acquisition and ***v* = 0.025** for long-term role order knowledge acquisition. These determine the size of the update at each timestep.

### Simulating long-term learning

First we ask whether the model can learn the potential roles of each word. To demonstrate this, we simplify the grammar with just two roles (Figure 5a). This could correspond to a “pivot grammar” (primitive two-element utterances in early childhood) (Braine, 1963) or an innate syntactic structure present in all children (Giusti & Gozzi, 2006). Words that tend to come first in this two-word phrase bind preferentially to the first role neuron. A separate set of words tend to follow, and associate with the second role neuron, thus establishing two primitive word classes.

**Figure 5.**
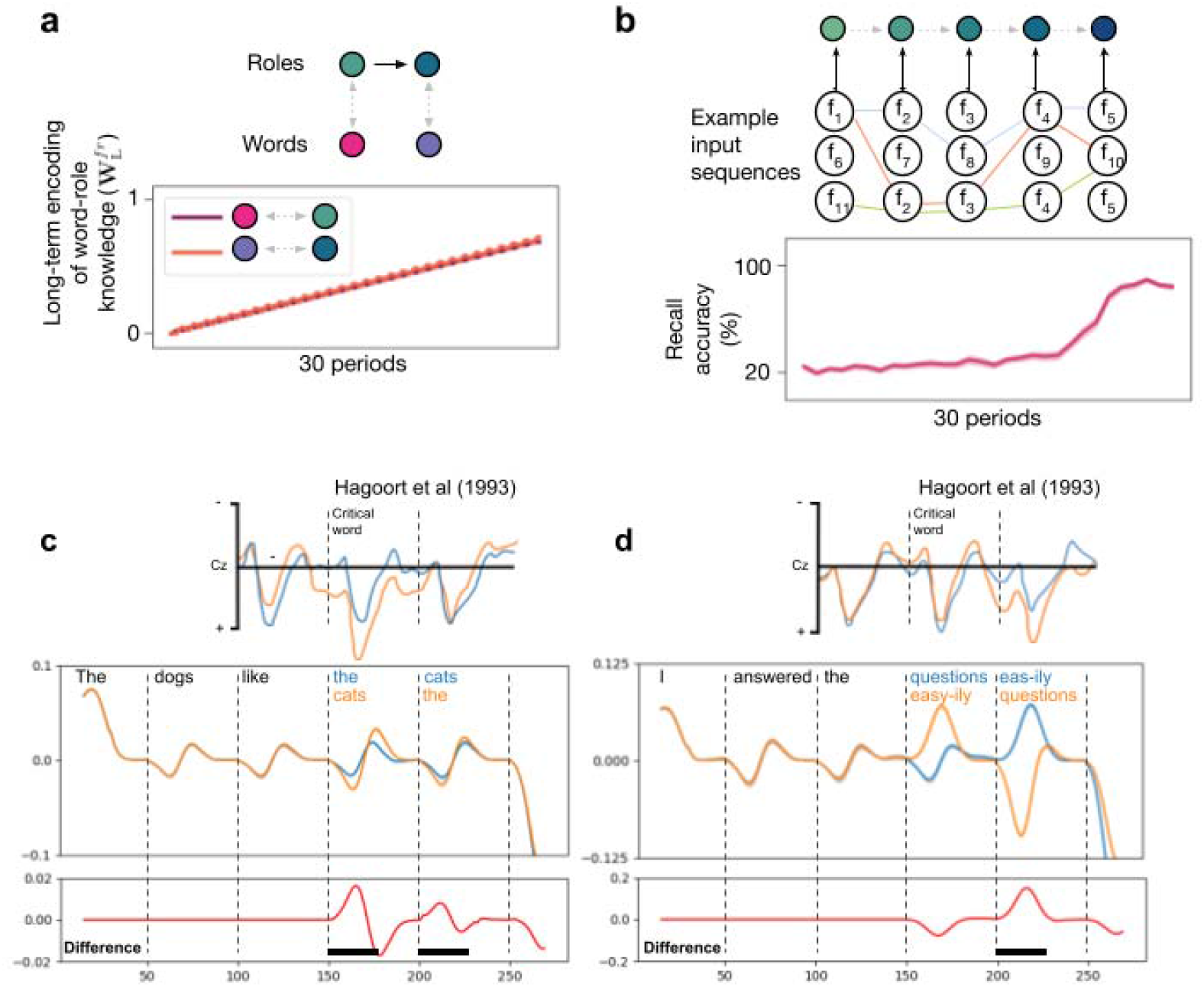
Long-term acquisition of word-class and syntactic knowledge. **a**, Simulation 13: Long-term acquisition of word class with pivot grammar. The role neurons parse only two-word sentences. The role neurons are activated sequentially, in tandem with the word input. This results in gradual learning between pairs of word and role neurons, representing long-term knowledge that some words appear in the first role, and others in the second. b, Simulation 14: Long-term acquisition of syntactic structure. The network receives randomly generated sentences with the same syntax over 30 periods (three possible words per role), and long-term knowledge between role neurons is learnt from the resulting order of activation of the role neurons. After each learning period recall accuracy is tested. Gray broken arrows indicate lack of predefined long-term knowledge. c, Simulation 15: Simulated event-related potentials (ERPs) from a syntactic vs non-syntactic sentence. Upper panel shows data from an ERP study, and the lower panel is the total neural activity in the model, passed through a delay, time derivative and smoothing filter. Larger neural responses seen after a syntactic violation resembles positive shifts in human ERPs (black bar). d, Simulation 16: Simulated ERP from correct vs incorrect affix conditions. Positive shift in total neural activity in the model matches positive shift observed in ERP study (black bar). The networks are presented in Figure A9. Error band is SEM.

Second we challenge the model to learn connections between role neurons. After word-to-role mappings have been established, new orderings of these words drive new role sequences (Figure 5b). Frequent role orders become embedded in the long-term role-to-role weights. Even branched role-to-role connections can be learned (Figure A10). This aligns well with theories that propose both learning and priming might arise from a single plasticity rule (F. Chang et al., 2000).

While word classes and role sequences can be learned independently, it remains unclear how they organically emerge together in a developing child, with overlapping time frames during language acquisition.

### Simulating Evoked Potentials

Evoked potentials from the electroencephalogram (EEG) have provided neural signatures of syntax for over 40 years (Clardy et al., 1979) in many languages (M K & Prema Rao, 2020), and remain a subject of active research (Glushko et al., 2022). The ERP is an amplified, filtered and averaged electric field from the brain recorded through the scalp in response to a stimulus. It is believed to encode a bulk signal dependent on neural firing rates across large areas of cortex (David et al., 2006; Pinotsis et al., 2014). Since the model’s response to incoming words varies dynamically, it could therefore generate physiological predictions. To simulate EEG data from a single trial, we took the sum of the squared activations of all the model’s neurons. We then applied a high-pass filter, smoothed the signal over time and added a delay (Schöbi et al., 2021).

We asked whether these simulated ERPs distinguished between conditions where the grammaticality of a sentence was manipulated. We presented the network with either grammatical sentences, or sequences where one of the words violated the grammatical rules, either because the word class was inappropriate, or because it had the inappropriate affix. We compared the mean evoked responses for the two violation types with the grammatical condition. Word class violations generated a positive deviation in ERP, while violation in permissible affixes resulted in a negative deviation in evoked potential difference. These resultant ERP changes qualitatively resemble human EEG studies (Figure 5c-d) (Hagoort et al., 1993).

## Discussion

We describe a neural model that encodes words in a sentence together with their syntactic roles, separating their structure from content. It binds words to their respective roles via a rapid Hebbian rule. Even though this model takes in and outputs sequences of words, the internal representation encodes words by tagging them with roles -- which dissociates from word order in some languages. This role-based representation offers potential for future models to manipulate or transform the stored words, for example by external control over role-neuron activation.

While associationist theories of human language syntax have been strongly rejected in the 1950s/1960s, our model does not just represent items in an ordered sequence, but rather, it provides a backbone where a structural representation (in the present case, a branching Markov network) can be mapped onto word sequences. Crucially, our structure-mapping mechanism permits neural networks to work not with word-based associative sequences but with abstract structures.

The model predicts sentence superiority, grammaticalisation errors, types of aphasia (Datta & Boulgouris, 2021), and priming. It allows sentence generation, and languages requiring affix defined roles. The model relies on two core principles. First, winner-takes-all dynamics allows symbol-like discrete representations to activate in structured sequences (Figure A2). Second, Markov-like transitions between role neurons maintain structure over the words. This new architecture solves an old problem: How does the brain encode words as well as their function in a sentence?

Unlike large language models, our architecture is simple and interpretable. It may also be more biologically plausible, as rapid Hebbian plasticity requires only synapse-local information, and may be realized in cortical synapses (Fiebig & Lansner, 2017). Few neural models can *generate* as well as encode syntax – transformer-based neural networks achieve this using biologically implausible artifices, such as scrolling buffers and positional encoding. Our model also explains why violations of syntax lead to a larger neural response in evoked potentials, a key neural signature of syntax (Clardy et al., 1979).

Lesions in the model are able to produce separable agrammatic and fluent aphasias, often seen in stroke patients – but which are unlikely to occur in large language models.

Lesion effects have been modeled in terms of stepwise syntactic operations (Grodzinsky, 2000; Vosse & Kempen, 2000). These have met with variable success, since a wide range of language deficits are possible after brain lesions (Hoffmann & Chen, 2013). Artificial neural networks are also good at simulating a spectrum of deficits, but the classic dissociation between fluent and nonfluent aphasias has been the most challenging (Y.-N. Chang & Lambon Ralph, 2020; Hillis et al., 1999; Roelofs, 2022). Our model has achieved a clear distinction in the fluent (preserved structure) vs nonfluent (non-preserved structure) lesions it produces.

The approach further allows us to model errors and response times. The explicit symbol-like behavior, where structural rules are separated from content, could allow our architecture to be applied to other sequential cognitive processes, such as logical reasoning or deduction. In contrast with approaches that link neural networks to pure symbolic systems (Mao et al., 2021; Smolensky, 1990) a sentence is encoded in synapses, rather than as a pattern of neural activity.

Previous models learn to produce sequences. However, they produce sequences of *tokens*, rather than a sequence of roles that bind to tokens. To implement syntactic rules, we need to process words sequentially, while associating each with a role that depends on context. Role activation does not follow a fixed sequence, since the roles are driven not just by the previously active role, but also by word classes, by word case and affixes, and can include branching structures. This requires qualitatively very different dynamics to previous models that simply remember sequences.

An alternative view to ours is that elements in a sentence may be bound into phrases arising from phase locking of neural oscillations (Hardy et al., 2023), or through timing and rhythm of phrases (Martin, 2020), but operational simulations are lacking. Moreover, no readout or manipulation mechanisms have been proposed yet for such systems (Segaert et al., 2018). Accounts where binding occurs through oscillatory synchrony (Senoussi et al., 2021) tend not to account for how the information will be used, such as in recall. It is possible that the oscillatory changes observed experimentally are a result of the synaptic changes we have proposed in our model, which might be simulated in a spiking version of our network.

Our directional plasticity requires that the role-to-role facilitation must depend on the postsynaptic role neuron also receiving inputs from word neurons. Is this plausible? We speculate that potentiation only arises if the postsynaptic action potential travels retrogradely up dendrites, but the dendrites must be concurrently activated by word axons that arrive proximally on the dendritic tree. Such 3-way plasticity gating is a strong prediction of our model.

The simple model presented has a number of important limitations. Presently, it does not reflect the hierarchical nature of sentence structures. This means that, to encode sentences like “the dog that chases the cat runs”, the current model needs to duplicate all its syntactic rules for the subordinate clause (Poletiek et al., 2021), and the model’s linear order will not explain rules of agreement (Everaert et al., 2015). Solving fully hierarchical grammar is beyond the scope of this paper, but we suggest there are ways in which this could proceed from the current model.

There are two major obstacles to implementing hierarchy in the present model: structure re-use, and position memory. The first obstacle, structure re-use, means that certain role structures (sometimes called motifs or templates (Snijders et al., 2009)), such as those that allow adding an adjective before an object, may be re-used within phrases or clauses. If there were a single neural representation for ‘adjective before object’, it will need to maintain separate associations with different adjectives, within different phrases. Rather than having multiple duplicate roles units, one solution is to use dynamic population partitioning (Figure A11) (Hayworth, 2012). The second obstacle, position memory, means that when a subordinate clause is being processed, the system needs to ‘hold active’ the position in the main clause, to keep track of the current position, so it can return there when the subordinate clause ends. One solution is that working memory sustains activity of the role neuron that introduces the clause, until the end of the clause. This sustained activity allows the parsing to return to the main clause in the correct state. However, this still limits the model’s ability to simultaneously assign the same word to two different roles, in the example “the little star is beside a big star”, and non-adjacent subject-verb agreement (e.g. “The <u>girls</u> from Paris <u>are</u> singing.”) (Adger, 2015) remains a key challenge to the model.

Even if our model were capable of parsing hierarchy, it would still fail to capture the abstract structure of mathematical languages, since it is ultimately an instance of a finite state machine.

We also do not model word recognition, effects of semantic context (Ralph et al., 2017), and do not address general variable binding that would allow semantic inference (Feldman, 2013). It also remains an open question whether learning rules, such as those proposed for learning word roles and role order, would be able to reliably acquire context-sensitive mappings in larger, more complex grammars. As our aim has been in demonstrating the biologically plausible nature of this simple model, we have focused on qualitative evidence, which limits the ability to measure model fit relative to other models. While it is possible to fit the model quantitatively to data, this would introduce additional free parameters.

A critical next step for our model is to demonstrate that a symbolic model like this could be scaled up to formulate blocks of text. One important piece of work needed to scale this network up is fine-tuning the curriculum and network parameters and structure. During learning, a mechanism may be required to create branches in the role structure. This is particularly important in the absence of hierarchical representation, which leads to more complex sets of possible sentence structures.

It is also unclear how the semantic-syntactic distinction is embodied in the brain. While neurons in superior temporal cortex are selective for the identity of auditory stimuli (Belin et al., 2000), neurons in prefrontal cortex encode abstract categorization of input (Antzoulatos & Miller, 2014; Freedman et al., 2003; Shima et al., 2007; Tanji et al., 2007) and might therefore represent word roles (Gwilliams et al., 2022). However, the dissociation is unclear in functional MRI, where both syntax and semantics activate a distributed network of dominant hemisphere areas (Rodd et al., 2015; Vigneau et al., 2006). One possible reason for this is that during any kind of syntactic and semantic language processing, both kinds of neural representation are requisite.

The strengths of our approach are its breadth, accounting for cognitive, psychophysical, neurophysiological and neurological findings, while remaining so simple as to be fully transparent.

## Conclusion

This simple neural model we have built presents a biologically plausible solution to the filler-role binding problem in language. It processes a sequence of words to produce a structured representation in working memory. We have shown that by separating syntactic rules from content, we can reproduce agrammatic and fluent aphasia, and can serialise a set of words into a grammatical sentence. While hierarchy, pronouns and agreement remain open challenges, the model presents a foundational first step. It forms a critical bridge between neural and symbolic processing, and forms the basis for more complex solutions to address the hierarchical nature of language.

## Appendix

### Supplementary Methods

#### Simulating simple grammar in working memory (Simulations 1, 2, 3, S1, S2, S3)

In simple recall comparing a syntactic sentence with a shuffled word-list, the model either receives a syntactic sentence or a shuffled word-list during the encoding phase. Each word is activated for 50 timesteps. Each repeat of a sentence or a word-list defines a single period. Then the model is prompted (i.e. with 5 steps of activation of the first role neuron) to recall the encoded sentence. In recall of sentences where there is switching, the model receives the first sentence for encoding and then is prompted to recall the first encoded sentence. Following which and without resetting, the model receives the second sentence for encoding (i.e. the switch) and it is then prompted to recall the second encoded sentence.

Recall accuracy is the proportion of input words that appear during the recall phase. A word is considered to have “appeared” when it has the strongest activation amongst all words of a timestep (*Argmax*). Recall accuracies are averaged over 200 trials with ±0.5 uniform noise in all connection weights. Human data on recall accuracy is adapted from Allen et al (2018) Figure 1a by averaging the recall accuracy at each serial position. Statistical significance is calculated using unpaired, two-tailed Student’s *t*-test.

Grammaticalization errors made by our model are measured as the proportion of recalled word pairs that were syntactic. The 8-word long sentence is shuffled randomly 30 times. Each of these 30 shuffled word-lists is encoded and recalled by the model over 200 trials with different random seed initialisation and ±0.5 uniform noise to all connections. One such word-list has been included in Figure 1b(iii) (“Up woke for cat tea the its late.”) Statistical significance is performed using Pearson’s Chi-square test on summed total syntactic and summed total non-syntactic word pairs in recall over the 200 trials of all 30 word-lists. The null hypothesis is that there is no difference between the observed count of syntactic word pairs versus the expected count of syntactic word pairs, which is the chance level seen in Figure 1b(v) (red dashed line). The expected count is one-eighth (7 syntactic word pairs out of all 56 possible pairs in a 8-word sentence) of total number of word pair appearances during recall. The proportion of grammaticalization errors in Jones and Farrell 2018 is adapted from their Figure 3. It is calculated as the average proportion of improvement in 6-token parsing over all 20 participants.

Simulation of soft constraint is achieved by varying ***k***_**L**_ of 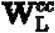 and 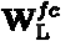 for connections “determiner” to “adjective” (***j*1**), “determiner” to “subject-noun” (***j*2**), “rich” to “subject-noun” (***k*1**) and “rich” to “adjective” (***k*2**). Heatmap is generated by repeating 2601 trials with ±0.25 uniform noise in all connections. Each trial involves defining the values of ***j*1, *J*2**,***k*1**, and ***k2***. Then, the network is encoded with the sentence “The rich men have money” and prompted to recall (i.e. parse). The network is then reset and encoded with the sentence “The rich have money” and similarly prompted to recall (i.e. parse). Each square in the heatmap is a single trial. Neighboring trials in the vertical direction differs by 0.01 for ***j*1** and ***j*2**, with ***j*1** increasing and ***j*2** decreasing from top to bottom. Neighboring trials in the horizontal direction differs by 0.01 for ***k*1** and ***k*2**, with ***k*1** decreasing and ***k*2** increasing from left to right.

Simulation of word class disambiguation uses the same constants as Simulation 1.

#### Languages with flexible word order requiring affixes (Simulations 4, 5, 6)

The model receives input sentences which are either words or pairs of words and affixes (50 timesteps for each word or pair) and is then prompted to recall. The lack of long-term role-to-role knowledge results in the need for stronger inhibition and activation between all neurons. Therefore deactivation of the preceding role neuron with an accompanying affix will require stronger inhibition between the role neurons, which has to be coupled with higher ceiling of activation to progress firing to the following role neuron. Likewise stronger activation between word and role neurons is needed to overcome the stable activation of the preceding role neuron accompanied with an affix. This too has to be balanced with stronger inhibition between word and role neurons. These parameter changes are reflected in Table A1.

#### Syntactic serialization, agrammatic and paragrammatic aphasias (Simulations 7, 8, 9)

The model receives the sentence (with each word taking 5 timesteps) or concurrent word inputs (over 20 timesteps) to be encoded and is then prompted to recall the syntactically structured sentence, leading to either control or aphasic output. During serialization, grammaticalization is achieved by including ***α*** inside the ***σ*** function. Closed-class words have lower ***k***_**L**_ in 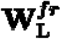, to ensure these words have connections more sensitive to noise in the network. The agrammatic lesion adds ±0.25 uniform noise to the rapid synaptic plasticity between role neurons 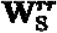. Intuitively, this makes the network less sure of the role of the next word in the sequence. The paragrammatic lesion adds ±0.005 uniform noise to long-term connections between word and role neurons, and with **λ**^***f*r**^ = 0. Intuitively, this means that while the role sequence is grammatical and grammatically appropriate words will be selected, the word will not be the encoded one.

Syntax errors were the number of non-syntactic word pairs in the output, including situations where more than one word had the highest activation at a timestep. Word errors were the number of words appearing in recall but were not in the input. Both agrammatic and paragrammatic aphasias produced by the model were repeated 50 trials each, with different random initialisations. Human data is taken from Table 1 of Caramazza et al (1981) with the agrammatic error counts being the mean of the Broca’s aphasia cases. Statistical significance for the model is calculated using unpaired two-tailed Student’s *t*-test.

Errors in the agrammatic simulation included dropping the closed-class word, stopping the sentence early, and skipping intermediate words.

#### Affix hopping (Simulation S4)

Affix hopping is the phenomenon where an affix that belongs early in a sequence of words occurs later in the sequence. For example, in the English sentence “the dogs were eating”, the participle ending “-ing” indicates continuous tense, but does not attach to the active verb “were”, which should indicate tense but here is modal.

Affix hopping can arise in the model because once a particular syntactic branch is chosen, there is no requirement for affix to connect to only the first role neuron in the branch. To demonstrate this, Simulation S4 adopts all parameters from Simulation 7 with the exception that concurrent word and affix (or semantic tag) inputs are run over 30 timesteps. The model receives either a sentence with simple active verb (e.g. “the dogs eat” with semantic tag for the role neuron of the active verb) or a sentence with a past participle (e.g. “the dogs were eat-ing” with a semantic tag for the auxiliary role neuron with strong connection to “were”. The affix neuron “-ing” is not activated during the encoding phase and is expected to spontaneously activate during the recall phase. “eat” in both sentences belongs to the same single word neuron (Figure A8).

#### Lexical and syntactic priming (Simulations 10, 11, 12, S5, S6, S7, S8)

In simulations of lexical and syntactic priming effects on comprehension, the model receives a pair of sentences to encode with no reset in between. For instance in a_1_b_1_, the model is encoded with priming sentence a_1_ and then target sentence b_1_, each over 50 timesteps. The model is then prompted to recall the target sentence. Recall accuracy follows protocol set out in Simulation 1. Each simulation is repeated over 50 trials with ±0.25 uniform noise for Simulations 10, S5 and S7, ±0.15 uniform noise for Simulations 11 and S8, and ±0.1 uniform noise for Simulation S6 in all connection weights. Response time is measured by the timesteps elapsed to reach the final word as the strongest item (*Argmax*). Response times are calculated only if 3 or more words in the target sentence are successfully retrieved. Unpaired, two-tailed Student’s *t*-test is used for comparing recall accuracy and Mann-Whitney U test for comparing response time.

To simulate Bock (1986)’s syntactic priming experiment in language production, the network needs to include neurons for agent and patient, which we term “semantic tags”. They behave just like affixes, in that they drive a given role neuron, and can be co-active with words. The agent and patient semantic tags drive the role neurons for objects, but in different syntactic branches – one corresponding to a sentence where the object is an agent, and another where the object is the patient. The subject role is not constrained to either role because, in English, the subject can be parsed without knowing which semantic role (agent or patient) it will take. Therefore the subject role neuron has a semantic tag representing either agent or patient.

First, the priming sentence is encoded with no semantic tag activation. Each word is presented in sequence for 10 timesteps, driving the role neurons down one of the two syntactic branches, and strengthening those role-to-role weights.

Second, to simulate a mental idea, the network has to first refresh its semantic representation as the subject and object in this mental idea is different from the nouns in the priming sentence. Therefore, we activate each noun in the mental idea simultaneously with its appropriate semantic tag (agent or patient). This results in each noun being associated to one of two possible object role neurons; thereby one of the nouns will be assigned to the object role in the primed syntactic pathway. In addition, a subject semantic tag is also activated with both nouns. We associate *both* nouns to the subject role as we want the network to choose the subject noun in the next step based on the priming sentence. Therefore three neurons are activated at a time as a triad (e.g. triad containing “birds” + patient semantic tag + subject semantic tag then triad containing “hunters” + agent semantic tag + subject semantic tag). The action word in the mental idea is also activated, in order to associate it with both the verb and participle roles. Through rapid synaptic plasticity, association of this new action word helps overwrite the network’s association to the previous action word in the priming sentence. Likewise, the network will also spontaneously choose whether the action word in the mental idea is a finite verb (active) or participle (passive) during the next step, based on the priming sentence. The action word does not require any semantic tag and therefore only one neuron is activated (i.e. “shot”). Each triad or neuron is activated for two timesteps, separated by 20 timesteps between triads or neuron to avoid cross-association. This cycle is repeated 15 times. The reason for using short repeated encoding of the semantic tags, is because the role neurons would otherwise tend to progress to the subsequent words. From this stage onwards, the rapid synaptic plasticity between role neurons is switched off (**λ**^**rr**^ = 0), just as for recall, but word-to-role plasticity (**λ**^***fr***^) is still needed to facilitate selection of the appropriate word in the sequence.

Third, the network has to form the sentence before utterance in the next step. It therefore receives a “bag of words” consisting of semantic contents (“birds”, “shot” and “hunters”) and the network spontaneously serializes the word content into a grammatical sentence in either the active or passive voice depending on initial priming sentence. The network receives two timesteps of the first word “the” and then 20 timesteps of simultaneous input of words “birds”, “shot” and “hunters” for syntactic serialization. This allows the network to encode a grammatical sentence from just semantic representations of the mental idea. A crucial dynamical process is also taking place during this step. Recall that we mentioned in the second step that both “birds” and “hunters” were activated with the subject semantic tag, but clearly only one can be the subject of the target sentence. Therefore at this stage, the network dynamics weakens the connection between the subject role neuron and the noun attached to the object role in the primed pathway. This is possible as the noun attached to the object role has its weight to the object role neuron strengthened from increased activation of the object role neuron in the destined branch as determined by the initial priming sentence. The Hebbian dynamics thus results in the weakening of the connection between this noun and the subject role neuron. For example, in the passive priming case, “hunters” is the object of the sentence; thus its weight to role neuron “r_8_” is strengthened while the network dynamics weakens its weight to the subject role neuron “r_2_”.

Fourth, during the utterance stage, the network is prompted to produce the target sentence describing the mental idea by activating the first role neuron for 5 timesteps, similar to a recall phase. This results in the words being activated in sequential order, either corresponding to the active or passive voice.

Fifth, the whole process is repeated 1000 trials for each priming condition (i.e. active or passive voice) with ±1.5 uniform noise in all Hebbian connection weights. The resultant utterance is counted as an active voice if the words appear in the order “agent, verb then patient” regardless of presence or absence of other words. Likewise, passive voice is counted if the order is “patient, participle then agent”. If both sequences are present, the first sequence to appear is counted in keeping with Bock’s original experimental protocol. Human subject data is obtained from Table 1 of Bock (1986).

#### Long-term knowledge acquisition (Simulations 13, 14, S9)

In long-term knowledge acquisition, the model is given either partial or complete lack of long-term knowledge (i.e. weights in 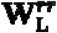 and 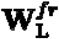 are zero) to simulate the naive learner. The network undergoes 30 learning periods and as before, during each period a sentence is encoded. In the case of long-term acquisition of word class knowledge using pivot grammar (Simulation 13),**W**_**S**_ is re-initialised to 0 at the end of every timestep to simulate a naive learner with no inhibitory connections at rest. In the case of long-term acquisition of role-role knowledge, **W**_**S**_ is re-initialised **−*k*** to at the end of every timestep. The learning rates for the long-term weights are much lower than for the short-term weights, but still follow the same Hebbian rules. Each neuronal activation lasts 50 timesteps. All other constants are unchanged from Simulation 1.

During long-term acquisition of word-to-role knowledge, the role neurons receive simultaneous activation with the word input to simulate an already-existing pivot grammar. Sequential activation of the two role neurons, as opposed to spontaneous activation of the second role neuron from the first role neuron, is required due to the naive network’s lack of inhibitory dynamics resulting in early activation of the second role neuron.

During long-term acquisition of role-to-role structure, the network receives a different randomly generated sentence for each period, but with the same syntax. Each role neuron has three possible words. Recall accuracy after each period is determined by running Simulation 1 (50 trials and ±0.5 uniform noise in all connection weights) on the network with frozen long-term weights and allowing for rapid synaptic plasticity.

#### Evoked potential (ERP) simulation (Simulation 15, 16)

To simulate an EEG for one trial, we apply a temporal derivative to the sum of squared activations of all the neurons. We then apply 30 timestep gaussian smoothing and a 15 timestep delay.

In Simulation 15, 120 trials were simulated. A sentence was presented to the network in the standard way. Half were syntactic (Determiner → Subject → Verb → Determiner → Object) and half were violations (Determiner → Subject → Verb → Object → Determiner). We used all combinations of 3 possible nouns and ±0.5 uniform noise was applied to all connections to generate within-condition variability. The EEG was averaged for each condition to obtain an ERP. Constants were kept the same as Simulation 1 with each word taking 50 timesteps. Human ERP data was adapted from Figure 7 of Hagoort et al (1993). Here, phrase structure violation coactivates multiple role neurons, pushing the neurons into a higher energy state, further from an attractor, increasing firing rate. This results in the positive ERPs at the labeled points.

In Simulation 16, 60 trials were simulated. A sentence was presented to the network with either the correct affix placement (Determiner → subject → auxiliary → verb → adverb with affix “-ily”) or incorrect affix placement (Determiner → subject → auxiliary → adjective with affix “-ily” → object). All combinations of 3 adjectives and adverbs were used and ±2 uniform noise was applied to all connections. Likewise, EEG was averaged for each condition to generate ERP. Constants were kept the same as Simulation 4. Human ERP data was adapted from Figure 1 of Hagoort et al (1993). Activation of the affix neuron “-ily” in the correct agreement condition results in increased total firing rates amongst all neurons. Consequently there is a positive ERP. The preceding negative ERP is also explained by the activation of “-ily” in the incorrect agreement condition.

The model does not include semantics or agreement rules, and so does not generate ERP components associated with violations of agreement.

**Figure A1.**
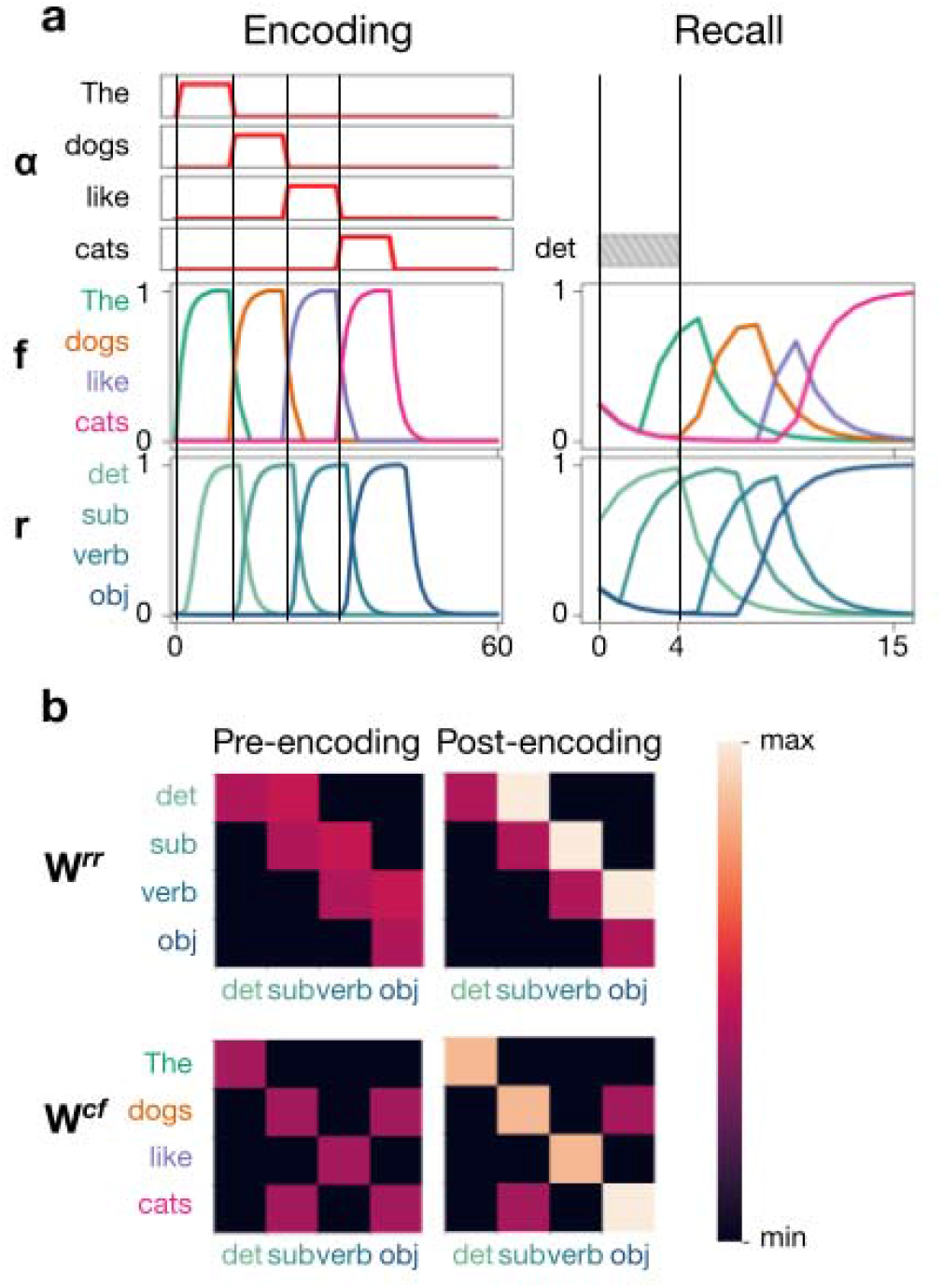
Simple walkthrough of encoding and recall of one sentence. a, Sensory input (**α**) results in activation of word neurons (**f**) and role neurons (**r**). Consequently, there is encoding through rapid synaptic plasticity of connections between neurons. This then allows recall of the sentence after brief activation of the first role neuron, the determiner. b, The sum of long-term and short-term synaptic weights before and after rapid synaptic plasticity. Role neurons: det=determiner, sub=subject, obj=object.

**Figure A2.**
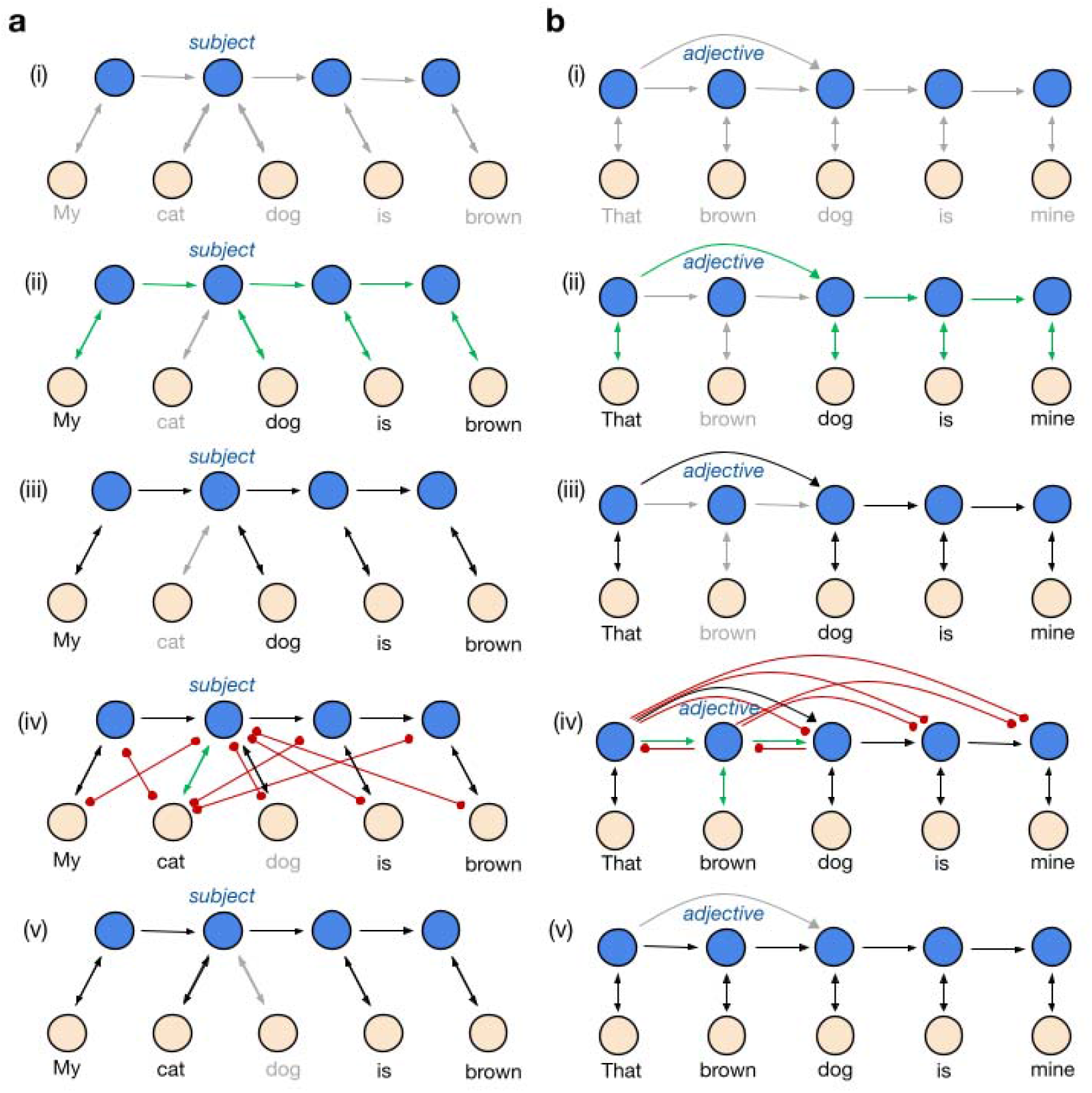
Graphical visualization of equations 5 and 6. a, Different words of the same class – e.g. both “cat” and “dog” – can be the subject of a sentence and therefore be connected to the subject role neuron. (i) Initial state of connection weights before any input. Gray arrows: long-term knowledge with no short-term rapid plasticity encoding. (ii) Reinforcement of some connections when presented with the first sentence “My dog is brown”. Green arrow: strengthening of connection. (iii) The resultant state after the first sentence, with “dog” associated with the subject role neuron. Black arrows: short-term rapid plasticity encoded. (iv) The concurrent reinforcement and weakening of connections when presented with the second sentence “My cat is brown”. (v) Final state of connection weights, with cat associated with the subject role. b, Distinct but overlapping sentence structures – the adjective “brown” may or may not be present in a sentence and therefore the adjective role neuron is skippable. (i) Initial state of connection weights before any input. (ii) Reinforcement of some connections when presented with the first sentence “That dog is mine”, with no adjective. (iii) The resultant state after the first sentence. (iv) The concurrent reinforcement and inhibition of connections when presented with the second sentence “That brown dog is mine”, recruiting the adjective role neuron. Red circle-ended arrow: weakening of connection. (v) Final state of connection weights, with the adjective neuron in the sequence.

**Figure A3.**
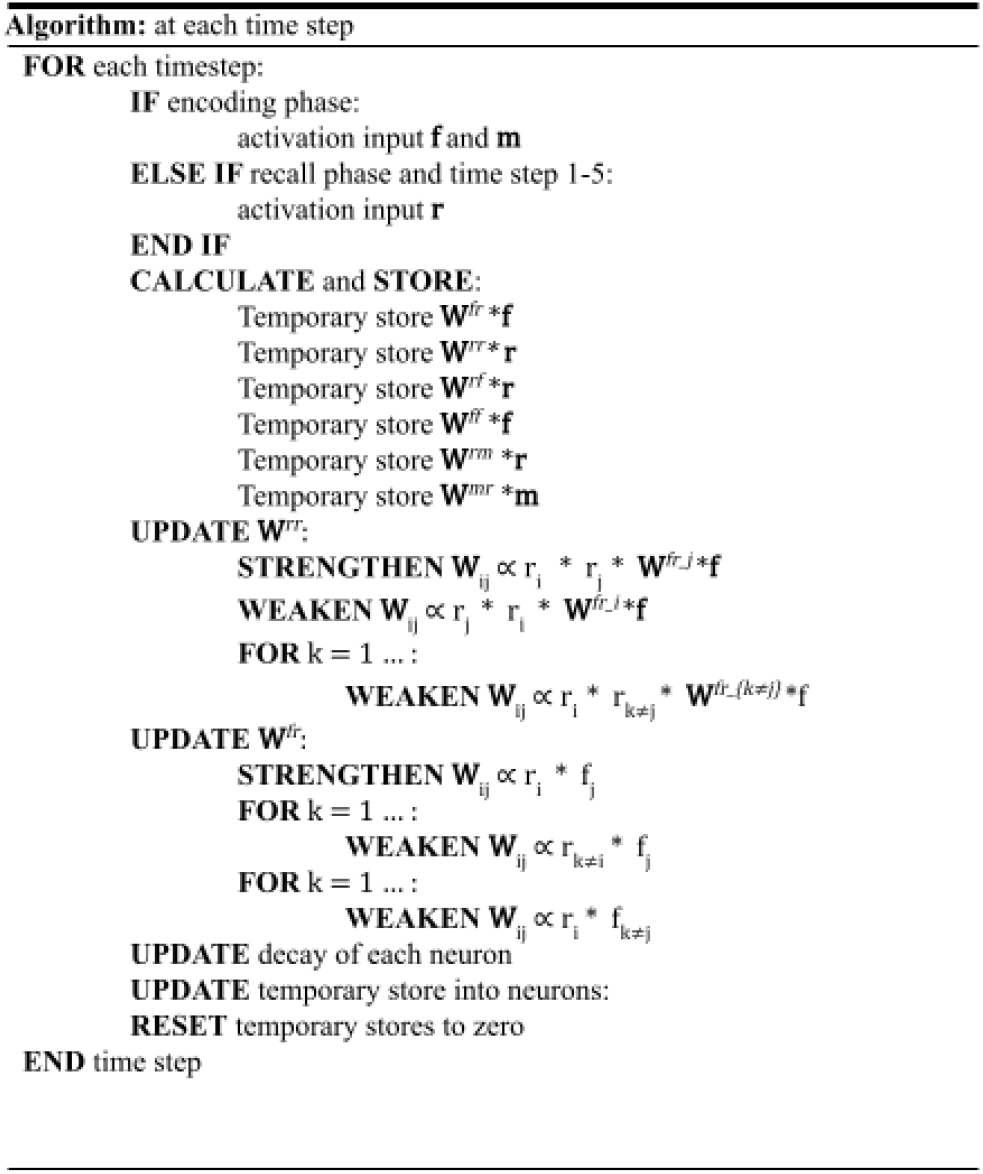
High-level pseudocode of the model at each timestep. The model receives sensory input to its neurons depending on if it is in the encoding or recall phase. The model then stores the output of all the neurons in a temporary memory. Rapid plasticity occurs as the short-term connection weights are updated. Each neuron then undergoes decay and their firing rate is then updated from values in the temporary memory store. The temporary memory store is finally reset to zero.

**Figure A4.**
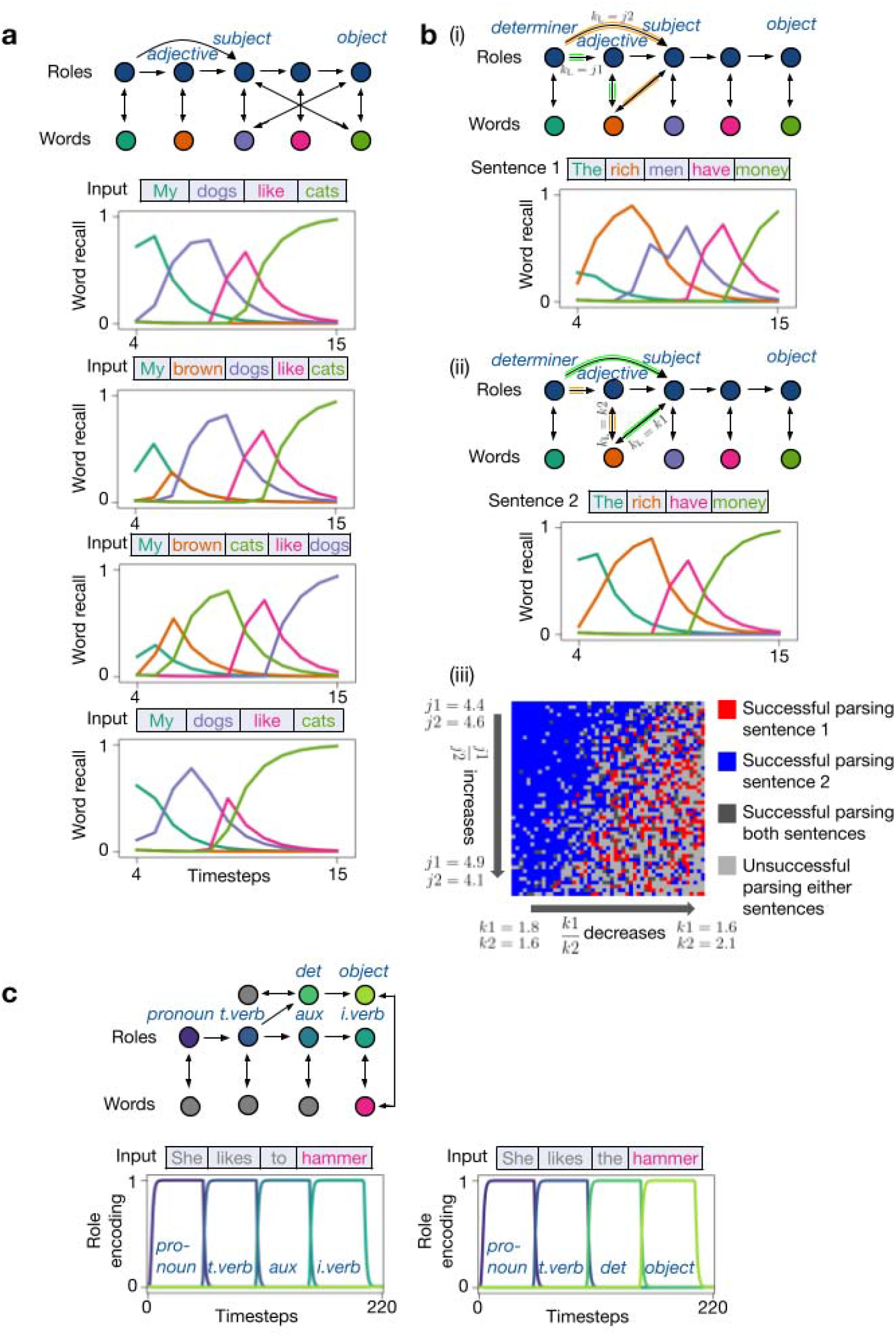
Additional simulations with simple grammar. a, Simulation S1: Both motifs from Simulations 2 and 3. Sequential rapid plasticity encoding and recall demonstrates the model’s ability to simultaneously handle both motifs. Notably in the final step from “My brown cats like dogs.” to “My dogs like cats.” there are both a switch from presence to absence of adjective and a switch between subject and object roles of the nouns. b, Simulation S2 of a garden-path sentence where “rich” could be an adjective or a subject. (i) “rich” preferentially binds to the adjective role neuron (***k*2 > *k*1**), which is stronger than the syntactic preference to skip the adjective role neuron (***j*2 > *j*1**). This results in parsing “rich” as an adjective. (ii) The word “rich” could also be parsed as a subject, but this leads to a garden path effect (failure to parse the sentence), unless the “rich”-”subject” synapse (***k*1**) is strengthened relative to “rich”-”adjective” (***k*2**), or the “determiner”-”subject” synapse (***j*1**) is strengthened relative to “determiner”-“adjective” (***j*2**). (iii) Varying the strengths of the syntactic and word-class constraints leads to different success rates in parsing the sentence. c, Simulation S3 of disambiguation of word class based on role sequence. Role neuron encoding is the activation of role neurons during the input of respective sentences. In “she likes to hammer”, activation of word “to” strongly activates the determiner role neuron which then activates the intransitive verb role therefore binding “hammer” to the intransitive verb role neuron. In “she likes the hammer”, the activation of “the” activates the determiner role neuron which prompts activation of the object role, therefore binding “hammer” to the object role neuron. This agrees with the “top-down” word class activation seen in MEG – that is the word role activated is one which fits the context.

**Figure A5.**
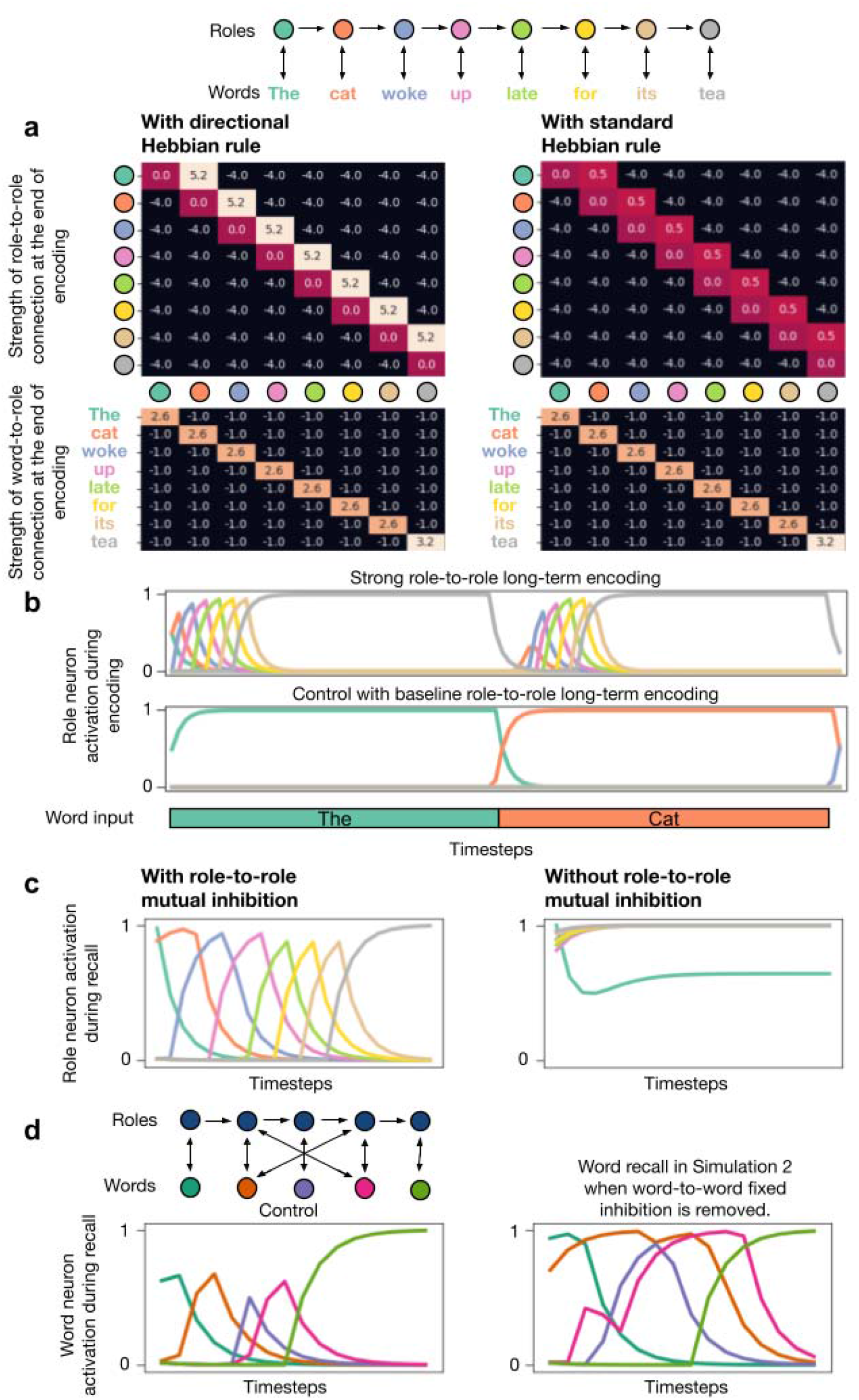
Effects of main architectural features. a, Without directional Hebbian rule, role-to-role encoding fails. There is no change to the word-to-role encoding. b, With strong long-term role-to-role encoding, there is sequential activation of all downstream role neurons during the input of each word disrupting the binding of one word to one role in the control scenario. c, Without role-to-role mutual inhibition, there is simultaneous activation of all role neurons during recall phase. d, Without fixed word-to-word inhibition, the model fails to recall the middle word (purple) of the sentence due to overwhelming activation of the two words sandwiching it as they both receive input from two role neurons each.

**Figure A6.**
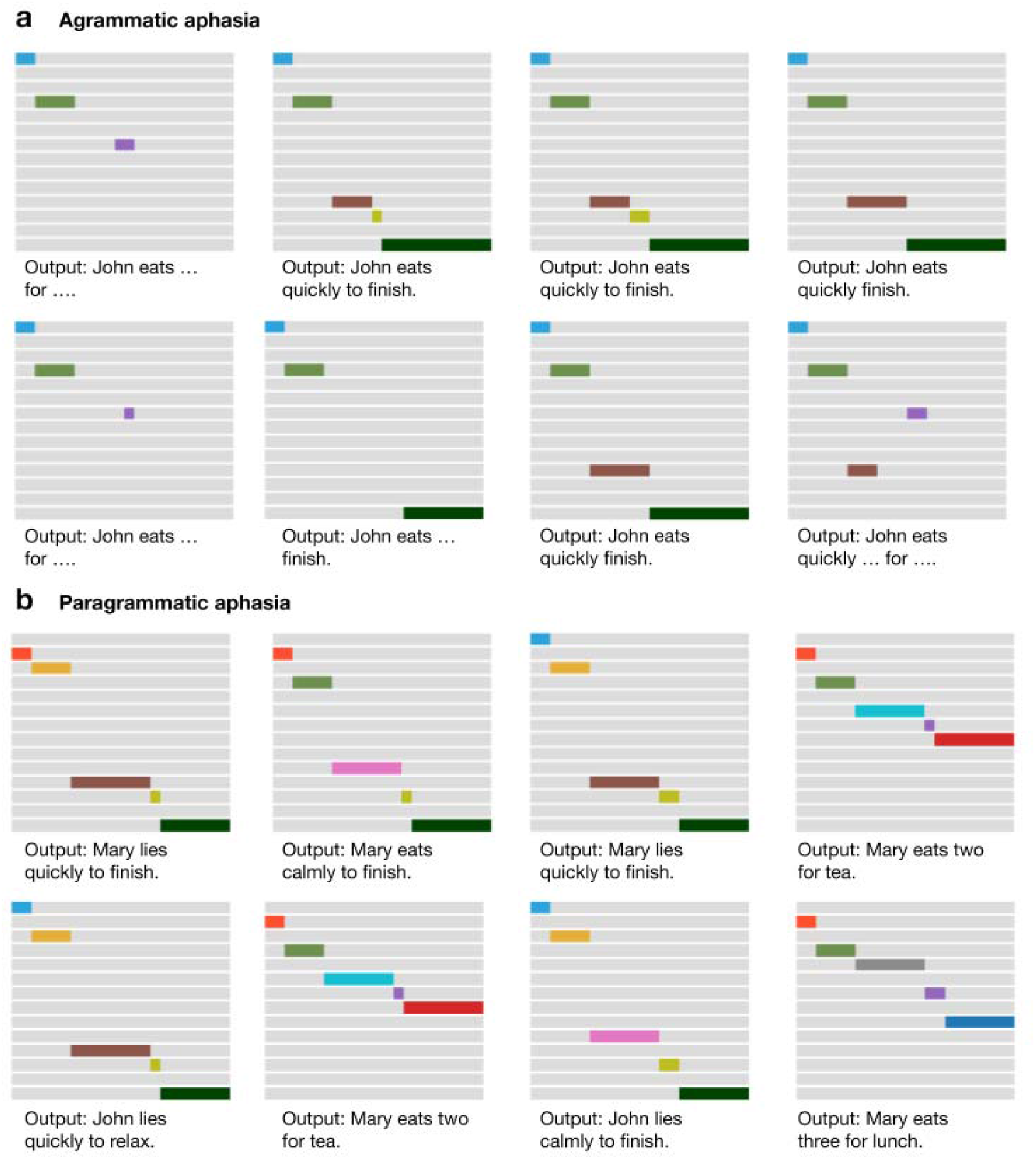
Network output with different random initialisations during simulation of agrammatic and paragrammatic aphasias. All network outputs are a result of the same hyperparameters but with different random initialisation seeds. a, Agrammatic outputs that are non-syntactic. ‘…’ represents more than one word having the same *Argmax* value. b, Paragrammatic outputs with characteristically well formed sentences but missing the underlying encoded message.

**Figure A7.**
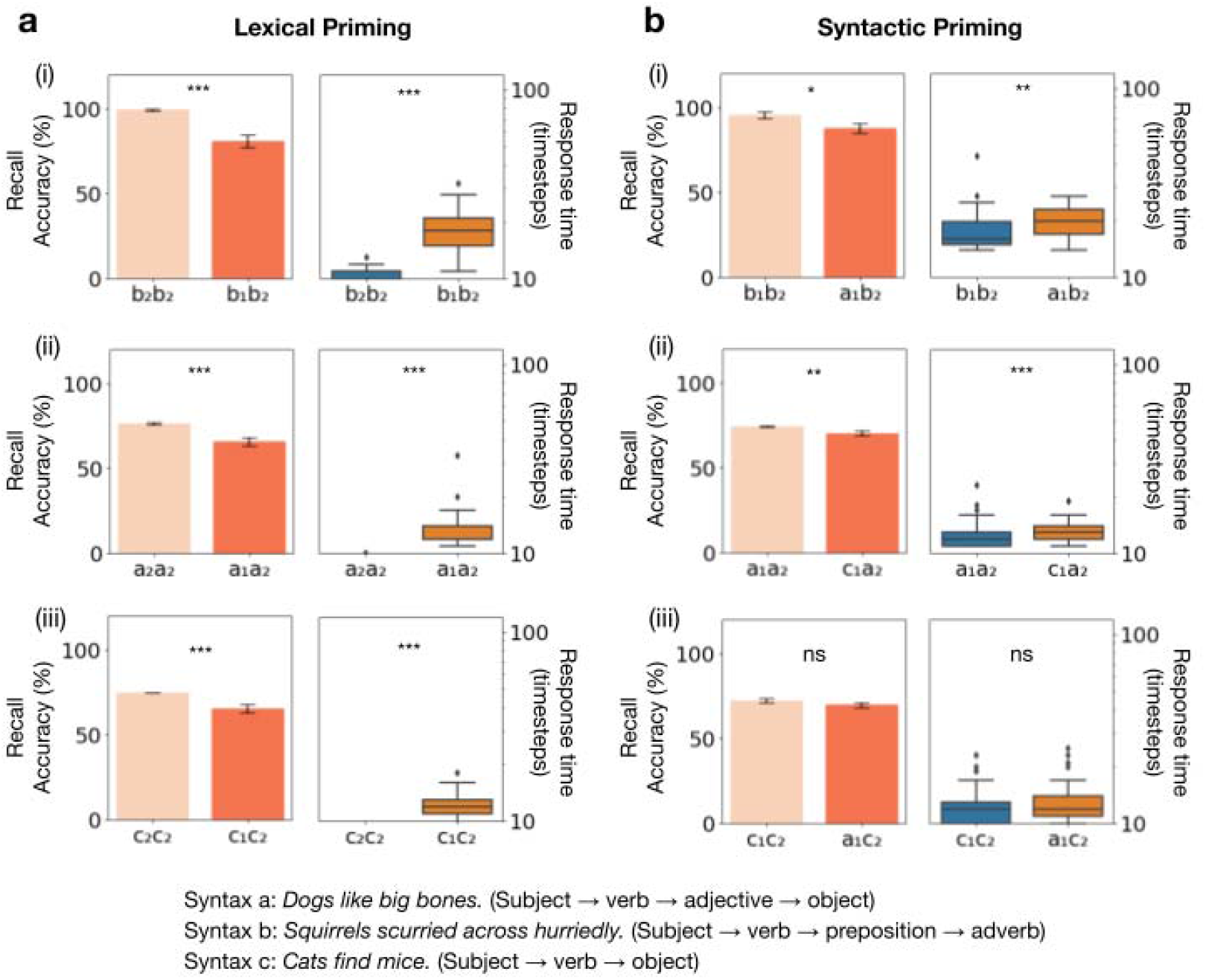
Results of lexical and syntactic priming during language comprehension. a, Recall accuracies and response times after lexical priming. (i) Simulation 10: Lexical priming of syntax b. (ii) Simulation S5: Lexical priming of syntax a. (iii) Simulation S7: Lexical priming of syntax c. b, Recall accuracies and response times following syntactic priming. (i) Simulation 11: Priming with either syntax a or b for target sentence of syntax b. (ii) Simulation S6: Priming with either syntax a or c for target sentence of syntax a. (iii) Simulation S8: Priming with either syntax a or c for target sentence of syntax c. Recall accuracies are with respect to the target sentence (i.e. second sentence in each pair). ns non-significant; *P-value<0.05; **P-value<0.01; ***P-value < 0.001, error bars are SEM. Box plot showing interquartile range.

**Figure A8.**
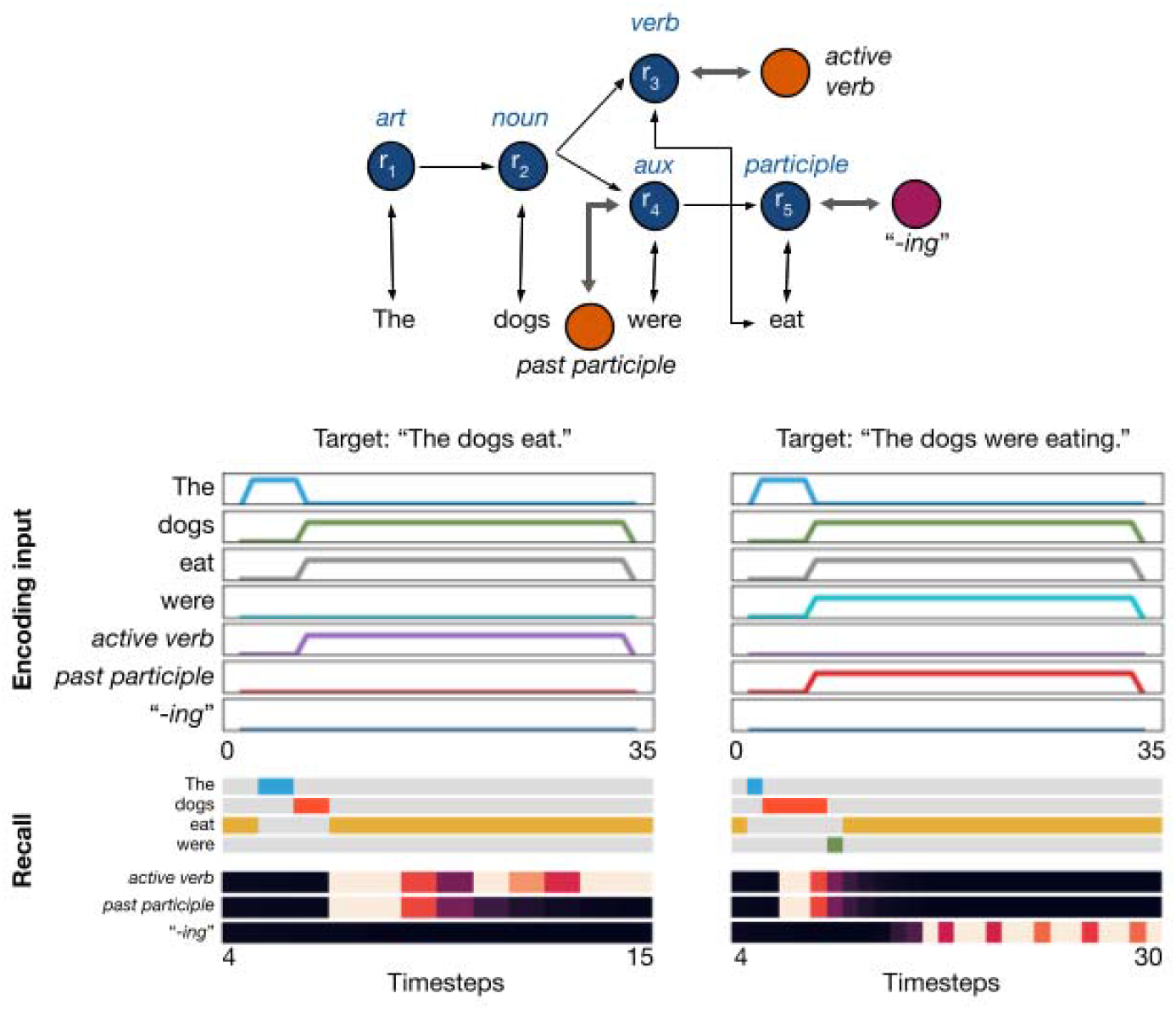
Affix hopping. In Simulation S4, the network is initiated with two possible tenses – simple present and past participle, each controlled by a semantic tag neuron which is functionally the same as an affix neuron. An affix neuron “-ing” is also tagged to the participle role neuron. The encoding input is a concurrent “bag of words” with respective semantic tags. Recall of the sentence in the past participle tense includes “-ing” activation with “eat”, simulating affix hopping.

**Figure A9.**
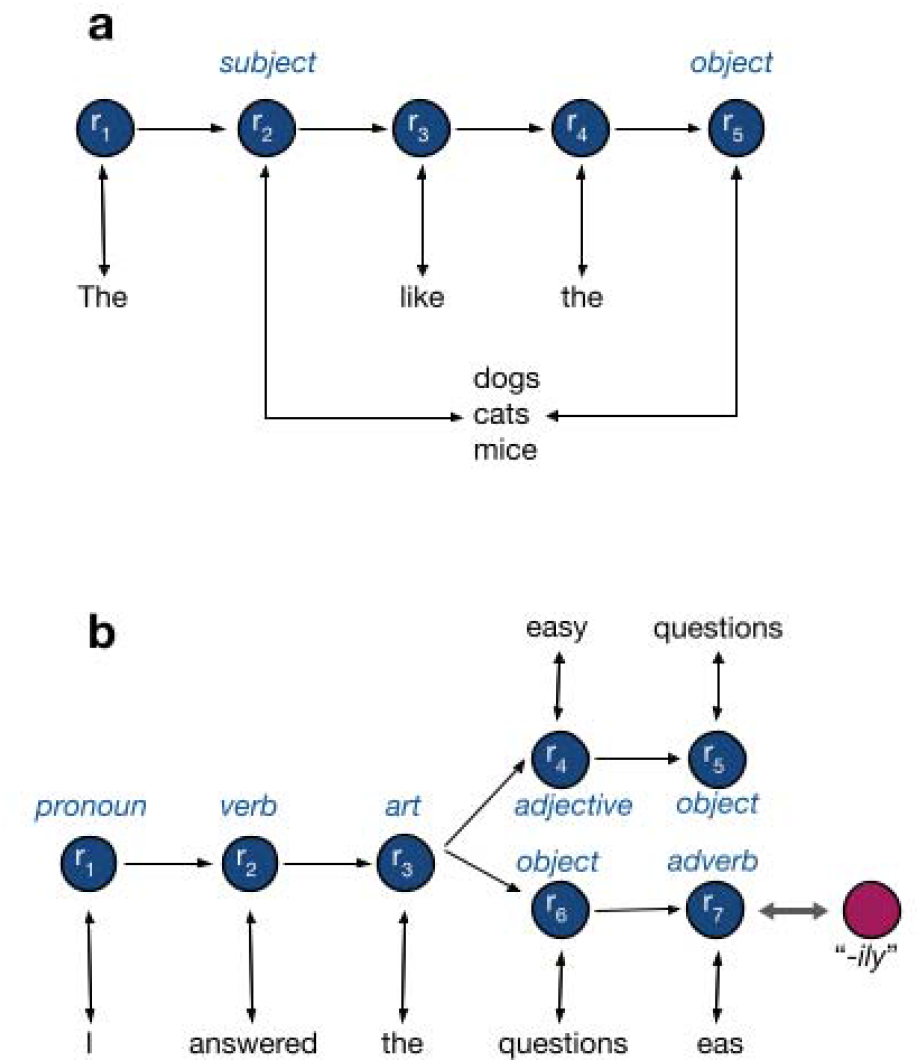
Networks of Simulations 15 and 16. a, A simple grammatical word order in the role neurons to demonstrate ERP during break in syntax. b, An affix neuron for “-ily” is bound to the adverb role neuron. In incorrect affix placement, “-ily” is activated with the adjective “easy”.

**Figure A10.**
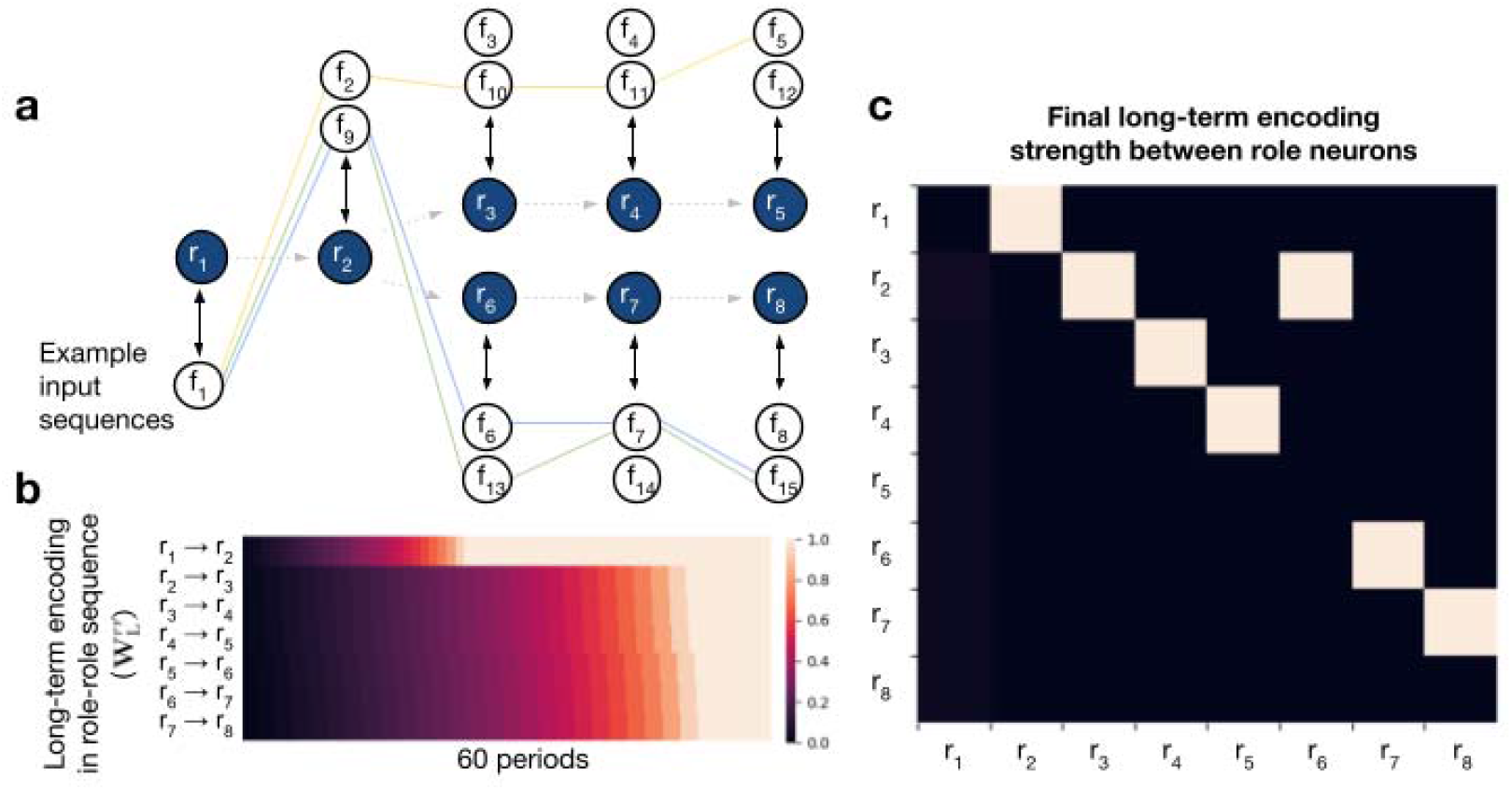
Long-term encoding of role-to-role knowledge with branching. a, Simulation S9: two possible syntactic structures amongst the role neurons can be learnt from the 15 available words. The model is fed randomly generated sentences from these 15 words. b, Strengthening of long-term role-to-role knowledge over 60 periods of interleaved sentences from each role sequence. c, The final long-term encoding weights of all role-to-role connections after the last period.

**Figure A11.**
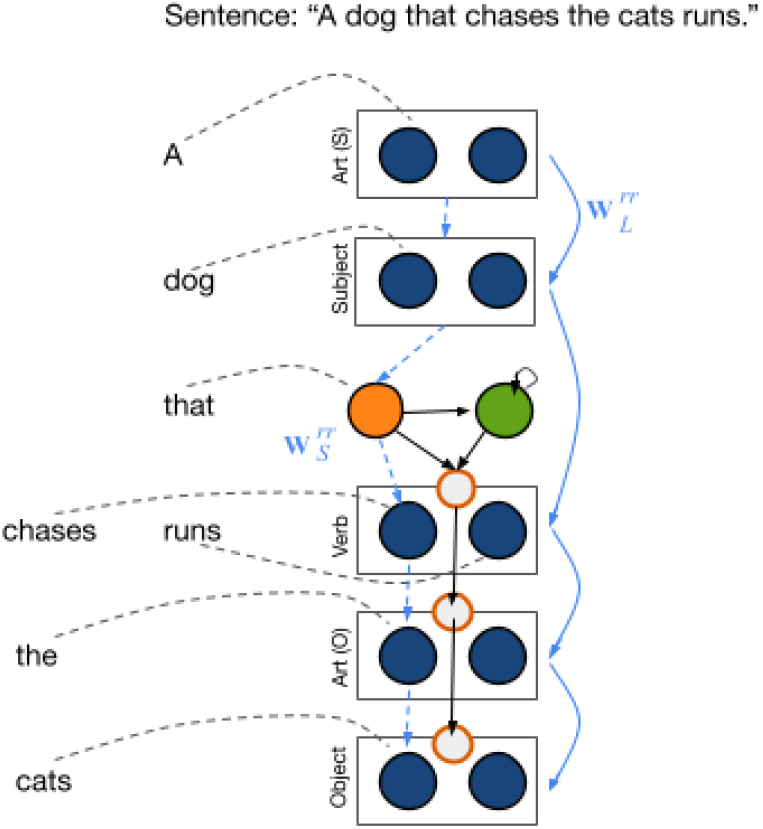
Potential mechanism to encode hierarchy. Two key steps are required to extend the model to encode hierarchical syntax structures. First, a mechanism must permit re-using previously used role neurons, to bind new contents. Second, a pointer to the main clause location must be maintained during the subordinate clause. For example, in the sentence “*A dog that chases the cats runs*”, there is an adjective clause “*that chases the cat*” and a main verb “*runs*” pertaining to the subject noun “*dog*”. The verb role neuron is used twice; once in the adjective clause as “*chases*” and once by the verb “*runs*”. We propose that this can be implemented by considering each role as a population which can be partitioned when needed (dark blue, two sub-groups of each role shown in figure). A pair of neurons indicating the start of an adjectival clause (orange) and a self-sustaining neuron (green) are activated on the word “*that*”. The green neuron actively maintains the state of the main clause, so that parsing can return to this point after the subordinate clause. The orange conjunctive role neuron activates an interneuron (grey with red border) which increases the lateral competitive inhibition within the verb role populations, such that two sub-groups compete and one wins. This effectively results in the dynamic splitting of the verb role population, allowing one of them to bind to “*chases*” while the other will later bind to “*runs*”. At the end of the adjective clause, the role sequence ends, permitting the green sustained neuron to drive the next role in the main clause, the main verb.

**Figure A12.**
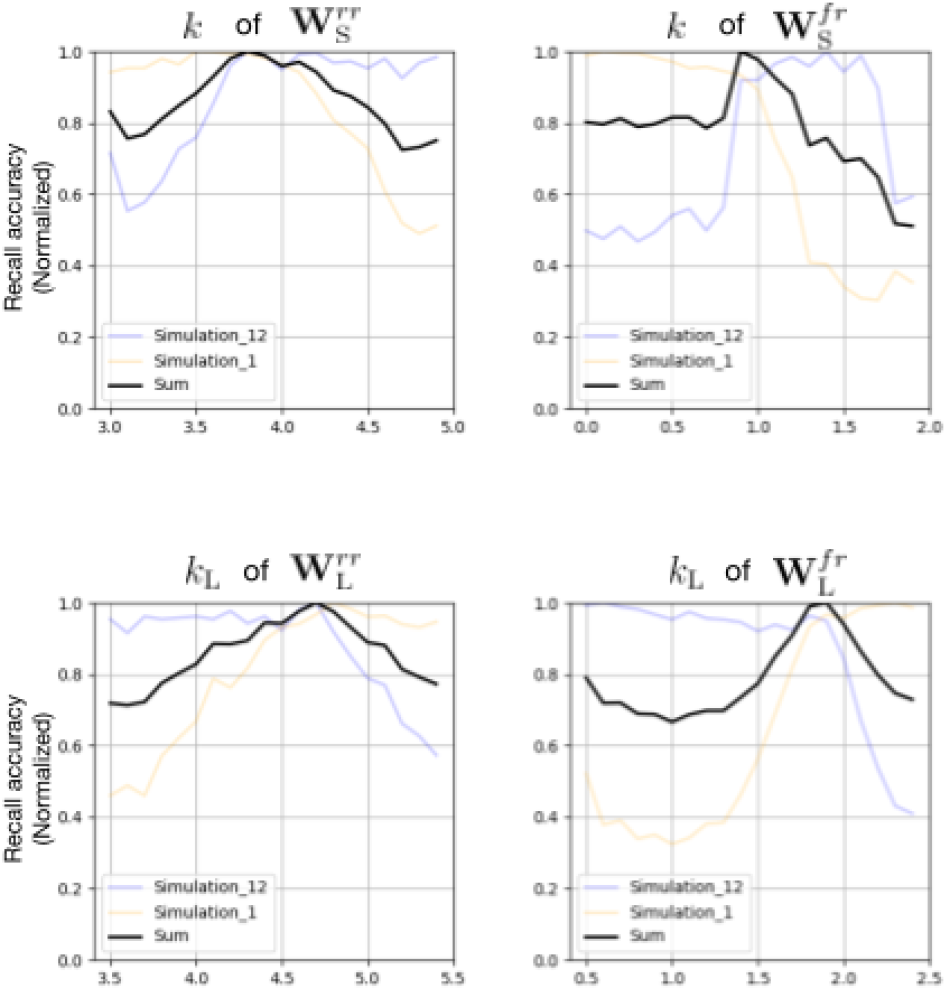
Parameter search. Accuracy of the model’s recall of a sentence is shown across a range of parameter values, for the role-to-role connections’ floor (top left) and long-term encoding scaling (bottom left), and the word-to-role connections’ floor (top right) and long-term encoding scaling (bottom right). Both Simulations 1 (simple sentence recall, orange line) and 12 (sentence generation with syntactic priming, blue line) are repeated with 100 different random seeds at 20 equally spaced values of each parameter (in the x-axis range shown). 0.5 and 0.1 noise were added to all connections for Simulation 1 and 12 respectively. Other simulations performed during parameter search have trends resembling either of the two simulations presented, and therefore are omitted. Peaks of each parameter search correspond approximately to the range we have provided in Table A1.

**Figure A13.**
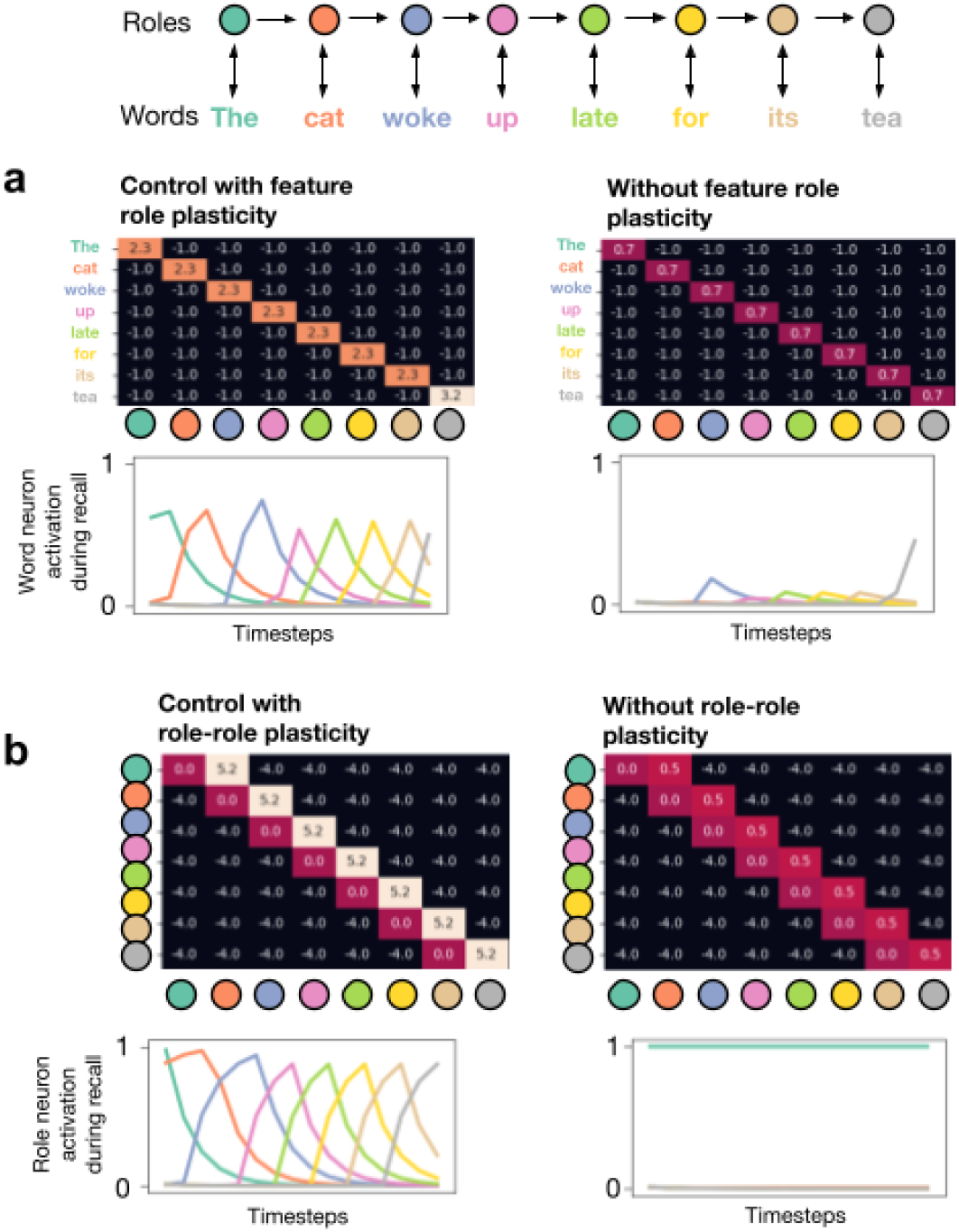
Complexity trade-off. a, comparison of word neuron activation during recall and final word-to-role connectivity (heatmaps) between Simulation 1 with feature role plasticity vs without feature role plasticity. The lack of strengthening in connectivity between words to roles results in poor activation of word neurons during recall. b, comparison of role neuron activation during recall and final role-to-role connectivity (heatmaps) between Simulation 1 with role-role plasticity vs without role-role plasticity. The lack of strengthening in connectivity between roles results in lack of sequential activation of role neurons during recall.

**Table A1.** Summary table of constants.

|  |  |  | Affixes | Simple<br>grammar | Priming<br>comprehensi<br>on | Priming<br>productio<br>n | Serializatio<br>n<br>aphasias |
| --- | --- | --- | --- | --- | --- | --- | --- |
| Simulations |  |  | 4,5,6 | 1,2,3,S1<br>,S2,S3 | 10,11,S5,S6,<br>S7,S8 | 12 | 7,8,9,S4 |
| f & m<br>Word and<br>affix<br>neurons | $\alpha$ | Input<br>strength | 1 (During encoding and with input*) | | | | |
|  |  |  | -1 (During encoding and without input†) |  |  |  |  |
|  |  |  | 0 (during recall) |  |  |  |  |

| r<br>Role<br>neurons | $\alpha$ | Input<br>strength | 0 (at encoding) | | | | |
| --- | --- | --- | --- | --- | --- | --- | --- |
|  |  |  | 1 to first role neuron on first 5 timesteps, then 0. (at recall) |  |  |  |  |
| $W_L^{fr}$<br>Long-term<br>word-role<br>knowledge | $k_L$<br>‡ | Weight<br>scaling | 3.7 [3.6-3.75] | 1.7 [1.7-1.75] | | | |
| $W_S^{fr}$<br>Short-term<br>word-role<br>memory | $h$ | Hebbian<br>baseline | 2 [2-2.25] | | | 6 [4-8]§ | |
| | $k$ | Minimum | 2 [1.95-2.05] | 1 | | | |
| | $\epsilon$ | Scale | 4 [2-4] | 1.5 [1.5-2] | | | |
| $W_L^{rr}$<br>Long-term<br>role-sequence<br>knowledge | $k_L$<br>¶ | Scale | 0 | 4.5 [4.5-4.55] | | | 8.25 [8.25-8.75] |
| $W_S^{rr}$<br>Short-term<br>role-sequence<br>memory | $\lambda^{rr}$ | Learning<br>rate | 50 [40-50]# | | | | |
| | $k$ | Minimum | 7.5 [7-7.5] | 4 [4-4.2] | | | |
| | $\epsilon$ | Scale | 4 [2-6] | 0.75 [0.5-0.75] | | | |
| $W^{mr}$<br>Affix-<br>role weights | | Scale | 6 [6-6.5] | | | 2 [1.5-2] | |

[Square bracket] indicates the range of values that the model is able to take with significant change to output, while keeping the other parameters at default values. This demonstrates the robustness of the parameters.
\*During serialization, **$\alpha = 0.8$** for words and affixes receiving input.
†During serialization, **$\alpha = -10$** for words and affixes not receiving input.
‡The closed-class words have **$k_L = 1.45$** for **$W_L^{fr}$** .
§The normalization term for Simulations 7, 8, 9 and 12 are higher due to more words being concurrently activated.
¶There is no long-term role-to-role knowledge during Simulations 4, 5 and 6. Simulations 7, 8 and 9 require stronger role-to-role long-term bias due to serialization, while Simulation 12 is exempt (which also has serialization) due to its priming phase.

